# The dynamics of introgression across an adaptive radiation: examining hybrid speciation and parallel adaptation in North American *Vitis*

**DOI:** 10.64898/2026.02.23.707523

**Authors:** Tianpeng Wang, Christopher J. Fiscus, Jacob B. Landis, Noé Cochetel, Abraham Morales-Cruz, Dario Cantu, Jonas Aguirre-Liguori, Brandon S. Gaut

## Abstract

Hybridization between species can contribute to parallel adaptive events and, in some cases, speciation. However, the prevalence of introgression is rarely measured across multiple species, meaning that it is rarely integrated with landscape-scale processes or connected to adaptive repeatability. Here, we analyzed whole-genome resequencing data from 639 accessions representing 48 *Vitis* species. *Vitis* is notable for the domesticated grapevine (*V. vinifera*), multiple economically important North American species, and as an example of a temperate adaptive radiation. Our dataset included population-level sampling for 19 species, from which we reconstructed individual- and species-level phylogenetic frameworks for the genus. The analyses uncovered widespread evidence of introgression, comprising ∼14% of the average *Vitis* genome. Introgression was associated with geographic distribution between species, and highly admixed individuals were more frequently found near ecological niche margins. Genetic analyses further indicated that previously recognized hybrid taxa, *V. x doaniana* and *V. x champinii*, likely represent hybrid swarms rather than distinct species. We also assessed patterns of adaptive repeatability across eight species with denser population sampling. For six of the eight species, most of the detected sweeps overlapped with sweeps in other species; on average, more recently diverged species shared more overlapping sweeps. Parallel sweep events were driven mostly by introgression, although a substantial fraction (∼40%) was attributed to selection on ancestral standing variation. By integrating population genomic data across many species, this study highlights the central role of shared genetic variation across an adaptive radiation.

## INTRODUCTION

Genomic data have yielded remarkable insights into the extent of ongoing and ancient hybridization between species (Nieto Feliner et al. 2020). Hybridization generally reduces genetic divergence between species or populations, but it can also reassort genetic variants into beneficial combinations that foster adaptation to new ecological niches (Marques et al. 2019). The benefits of genetic reassortment may partially explain highly reticulate relationships in adaptive radiations like those of *Heliconius* butterflies (Edelman et al. 2019) and African cichlids (Meier et al. 2017). The historical importance of hybridization is also reflected by the proportion of extant genomes that owe their origin to introgression, ranging from ∼2% in cultivated sunflower (Hübner et al. 2019) and humans (Prüfer et al. 2014) to nearly 50% in other cases [reviewed in (Schumer et al. 2018)].

One potential outcome of hybridization is speciation. Hybrid speciation occurs when a cross (or crosses) between two parental species create a distinct hybrid that grows in a novel ecological niche. Hybridization regularly drives the formation of allopolyploid species (Soltis and Soltis 2009), but homoploid speciation – where the same ploidy level is maintained – can also occur. A classic example is the sunflower species *Helianthus anomalus*, which is reproductively isolated from its two parental species and inhabits a unique niche in sand dunes (Rieseberg 2000). Historically, there has been some debate over how often homoploid hybrid speciation occurs, but the answer depends partly on the definition of a hybrid species. Schumer et al.(Schumer et al. 2014) have argued that the hybridization event must contribute to (or cause) reproductive isolation to fulfill the definition of a hybrid species. However, since hybrid speciation events are usually historical, it is difficult to know whether hybridization led to immediate reproductive barriers or whether species gained reproductive isolation gradually over time. The requirement of reproductive isolation is also too stringent for plants, since many plant species are interfertile. Accordingly, others have argued for alternative definitions - i.e., that a hybrid species is established, persistent, and morphologically or ecologically distinct from its parents (Nieto Feliner et al. 2017; Schumer et al. 2018; Baumgartner et al. 2020). In theory, the persistence (i.e., age) and genomic history of putative hybrid species can be elucidated by population genomic analyses (Schumer et al. 2014), but analyses of hybrid taxa are uncommon.

Hybridization and introgression can contribute to another major but under-characterized phenomenon, which is adaptive repeatability. The study of repeatability focuses on a few major questions (Bohutínská and Peichel 2024; Chaturvedi et al. 2025). The first is whether adaptation occurs in the same genes across different species; that is, is there evidence of repeatability in the adaptive process? If there is, it suggests that solutions to adaptive challenges are limited genetically. A second question is the source, or mechanism, of repeatability. Repeatability between two lineages may to be due to three mechanisms (Bohutínská and Peichel 2024; Chaturvedi et al. 2025): i) *de novo* adaptive mutations in separate lineages, ii) selection on variation that segregated in the common ancestor of the two lineages or iii) adaptive introgression between lineages (Chaturvedi et al. 2025). The relative frequency of these three mechanisms is an open question, but its address impacts our understanding about the dynamics of adaptation. It has also been hypothesized that repeatability is a function of divergence time (Bohutínská and Peichel 2024) - i.e., recently diverged lineages are expected to exhibit more repeatability, due to either more recent common ancestry or more frequent gene flow. Addressing questions about repeatability requires population genomic sampling across many species, both to detect shared selected sweeps and also to assess its mechanisms.

Here we study genomic diversity, species relationships, adaptation and hybridization across the genus *Vitis*. The genus contains ∼75 species of perennial vines, with three belonging to subgenus *Muscadinia* (2*n* = 40) (Buck and Worthington 2022) and the rest to subgenus *Vitis* (2*n* = 38). Subgenus *Vitis* is economically crucial; domesticated grapevine (*V. vinifera*) has the highest total production value among fruit crops (FAO, 2016) (Anon)^,(Vivier and Pretorius 2002)^, and 80% or more of global viticulture relies on rootstocks derived from wild *Vitis* species (Ollat et al. 2016). Species of subgenus *Vitis* are distributed throughout the temperate Northern Hemisphere, with roughly half native to North America, half to eastern Asia, and one (*V. vinifera*) to Eurasia (Vitaceae). The subgenus is also notable because it represents a temperate adaptive radiation from close tropical relatives (Liu et al. 2016), making it a plant model for understanding morphological diversity (Ma et al. 2016; Baumgartner et al. 2020) and phylogenetic diversification (Jaillon et al. 2007; Goremykin et al. 2009; Wan et al. 2013; Liu et al. 2016; Wen et al. 2018; Talavera et al. 2023). The North American species are particularly important, both because they are the source of all common rootstocks (derived from species such as *V. riparia*, *V. rupestris*, and *V. berlandieri*, to name just a few) (Chen et al. 2024) and because phylogenetic treatments of the genus suggest that North America was the center of species diversification (Talavera et al. 2023). There is also evidence for reticulate histories among species (Zecca et al. 2019; Morales-Cruz et al. 2021; Nie et al. 2023), likely reflecting the fact that all subgenus *Vitis* species are interfertile (Moore et al. 2016). Finally, the genus contains taxa of recognized hybrid origin (Heinitz et al. 2019; Zecca et al. 2019), offering the opportunity to study putative hybrid speciation events.

We have amassed a whole-genome resequencing (WGR) dataset of >600 accessions representing 48 *Vitis* species, with an emphasis on North America. The primary difference between our data and previous large *Vitis* WGR datasets (Liang et al. 2019a; Dong et al. 2023; Guo et al. 2025; Marroni et al.) is that previous work emphasized cultivated grapevine (*V. vinifera* ssp. *vinifera*) and its wild progenitor (*V. vinifera* ssp. *sylvestris*) without sufficient population genomic sampling of wild relatives to glean insight into the processes that have shaped their adaptive radiation. Our dataset includes population genomic sampling for 17 North American species, including two putative hybrid species (*V. x doaniana* and *V. x champinii*). We use this large dataset to address four sets of questions. First, what is the extent of admixture and hybridization across species, and what can we glean about phylogenetic relationships given the potential for reticulate relationships? Second, given two putative hybrid species in the genus, what do population genomic data reveal about the timing and dynamics of their formation? This investigation is not only crucial to characterize the history of the taxa but also to provide insights into the broader, and sometimes controversial, field of hybrid speciation (Nieto Feliner et al. 2017; Schumer et al. 2018). The third set of questions centers on repeatability. How often are adaptive solutions, as measured by selective sweeps, repeated across species? What is the relative contribution of *de novo* mutations, segregating variation and adaptive introgression to repeatability? And, does repeatability scale with time? By taking a multi-species approach, our work illustrates the remarkable adaptive impact of introgression during an adaptive radiation.

## RESULTS

### Complex admixture signals

We called variants for 639 whole-genome sequenced samples belonging to 48 *Vitis* species and 10 outgroup species within the family Vitaceae against the *V. arizonica* v.2.0 genome assembly (Morales-Cruz et al. 2023). The dataset contained 501 samples from 25 North American species that were sampled in the wild, planted as cuttings, and maintained as a living collection at the University of California, Davis. It also included previously published samples from Asian *Vitis* taxa (30 species, 53 samples) (**Table S1**) and a subset of available data for domesticated *V. vinifera* (ssp. *vinifera*: 44 samples) and its wild progenitor (ssp. *sylvestris*: 39 samples) across their geographic range. After mapping and filtering sites for missing data and poor quality genotype calls, the final dataset consisted of 4,435,884 biallelic SNPs with < 5% missing calls and a minor allele frequency ≥0.01 across the entire dataset.

Previous work has shown that the phylogenetics of *Vitis* is complex, likely due to introgression that obscures true species relationships (Nie et al. 2023). We began our analyses by drafting a phylogenetic tree and examining patterns of ancestry across all accessions. For the latter, we pruned the SNP dataset for linkage disequilibrium (LD) and then inferred ancestry profiles using NGSAdmix (Skotte et al. 2013). An “optimal” number of K = 13 clusters was selected for the 48 species based on log-likelihood and ΔK statistics (**Fig. 1; Fig. S1-2**). We paired the admixture analysis with the draft phylogenetic tree based on the pruned SNP set **(Fig. 1)**, uncovering evidence for widespread admixture. Some admixture was due to shared ancestry with domesticated *V. vinifera* (Xiao et al. 2023). For example, nearly half of the *V. californica* samples were admixed with domesticated germplasm, consistent with documented hybridization events between these species (Dangl et al. 2015). Another observation was that many *V. arizonica* individuals had ancestry contributions from either *V. rupestris* or *V. girdiana,* with some individuals appearing as 50:50 hybrids. Finally, the mixed ancestry of the two putative hybrid species (*V. x doaniana* and *V. x champinii*) was supported by two separate clades of *V. x doaniana*, reflecting groups with different majority ancestry from one of the two putative parental species (*V. mustangenesis* and *V. acerifolia*) (**Fig. 1**). There were also two distinct sets of *V.* x *champinii* samples; one had the commonly accepted parentage (*V. mustangensis* and *V. rupestris*) but the second appeared to have separate origins, with *V. monticola* as one parent. The accuracy of ancestry assignments are influenced by factors such as genetic drift and linked selection, but these results nonetheless suggest widespread introgression among *Vitis* species.

**Figure 1.**
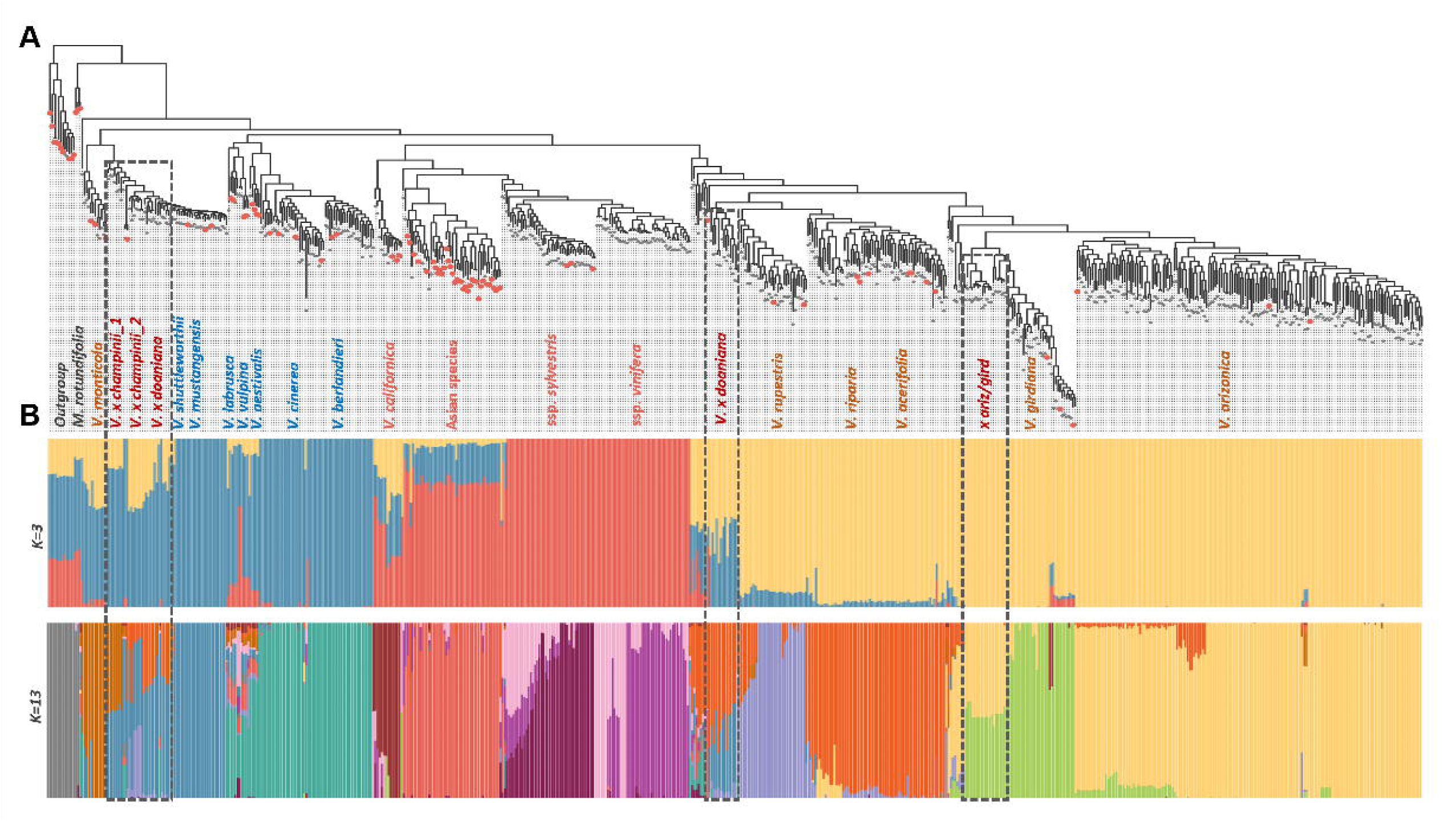
Phylogenomic reconstruction and population structure of 639 accessions. **(A)** The maximum Likelihood phylogeny of accessions inferred from genome-wide SNPs. Tips in red indicate the three representative non-hybrid individuals selected per species for species-level phylogenetic inference. Grey rectangles highlight key hybrid lineages (*V. x champinii*, *V. x doaniana,* and a *V. arizonica* / *V. girdiana* complex). **(B)** Model-based ancestry inference across the 639 accessions shown for K = 3 and K = 13. Each vertical bar represents one individual and colors indicate estimated ancestry proportions from the inferred K genetic clusters. Individuals are ordered based on their phylogenetic position. Complete admixture plots for K=2 to 15 are provided in Fig. S2.

### Species phylogeny and divergence times

Extensive admixture and reticulate evolution among *Vitis* accessions poses challenges for inferring a species phylogeny. However, investigating introgression and adaptive repeatability requires a phylogenetic framework, ideally with divergence time estimates. We therefore inferred species-level phylogenies, starting by removing samples with substantial ancestry admixture (>20%) or that did not cluster as expected with other members of their assigned species (**Fig. 1**). In total, 140 samples (22% of the dataset) fit one or both of these criteria and were excluded from further phylogenetic and population genomic analyses (**Fig. S3**, **Tab. S2**). Note that the removal of these samples is likely to make our population genomic inferences of introgression to be conservative (see below).

To infer the species phylogeny and estimate divergence times, we randomly selected up to three samples per species, producing a dataset of 92 samples across 48 species (**Fig. 1**). With these data we first constructed phylogenetic trees with RaxML and SVDQuartets (**Fig. 2A & Fig. S4**), using LD-pruned SNP genotypes. The two trees had highly similar topologies (Generalized Robinson-Foulds matrix distance = 0.19) but differed in at least one major respect: whether the monophyletic Asian clade was nested within the North American clade (**Fig. S4**). We also constructed trees based on K-mers to generate a reference-free cladogram and also on assembled chloroplast haplotypes; the K-mer tree had several rearrangements (R-F distance = 0.24 from both the RaxML and SVDQuartets trees; **Fig. S5**) but recovered the major clades from the SNP-based trees. In contrast, the plastid-based tree was highly discordant to the other trees (R-F distance from 0.44 to 0.48 to the three other trees; **Fig. S6**), as has been noted previously (Talavera et al. 2023).

**Figure 2.**
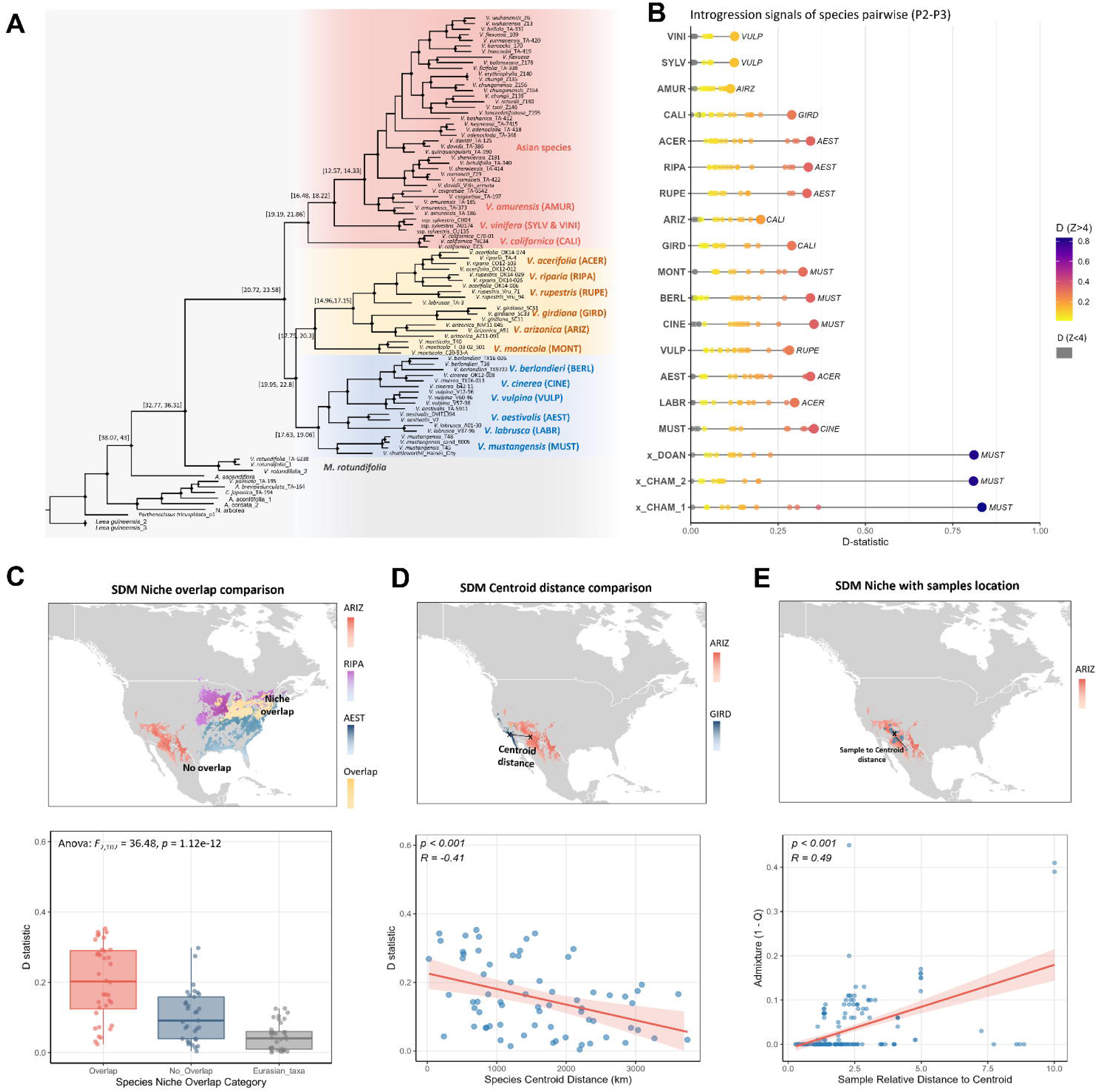
Phylogenetic history, introgression signals, and geographic correlates of gene flow among North American *Vitis*. **(A)** The SVDquartets phylogeny of non-hybrid *Vitis* with branch lengths estimated by maximum likelihood. Node ages were estimated with BEAST2 under a relaxed molecular clock (Fig. S7). Numbers at nodes indicate divergence-time estimates of 95% highest posterior density (HPD) intervals. **(B)** Genome-wide introgression signals from pairwise D-statistics for each P2 species (rows), with points representing various P3 species comparisons. Points are colored by the D-statistics; grey if non-significant (Z < 4) and following a gradient for significance *P*-value. The P3 pairing of the maximum D-statistic is labeled for each row. The top three species are Eurasian. **(C)** Top: Example species distribution models (SDMs) for *V. arizonica* (ARIZ), *V. riparia* (RIPA) and *V. aestivalis* (AEST) illustrating present-day habitat suitability and niche overlap. Bottom: Boxplots summarizing D-statistics for all species pairs (excluding hybrid species *V. x doaniana*, *V. x champinii*) that are grouped by SDM niche-overlap category: overlapping North American species pair, non-overlapping North American species pair, and other Eurasian *Vitis* species. **(D)** Top: Example SDM niche centroid-distance comparison for ARIZ and *V. girdiana* (GIRD). Bottom: A scatter plot summarizing the relationship between pairwise species D-statistics and centroid distance across all tested species pairs. **(E)** Top: Example map shows SDM-predicted niche for ARIZ. Bottom: The scatter plot summarizes the relationshipm across all individuals and species, between an individual’s admixture level and its location relative to the normalized distance to its SDM centroid. Individual admixture is quantified as 1 − Q, where Q is the individual’s maximum ancestry proportion from population structure at K = 13. Full SDM outputs for all tested North American *Vitis* species are provided in Supplementary Fig. 11.

Taken together, the phylogenies based on genome-wide nuclear data reveal at least four notable features (**Fig. 2A**). First, the *Vitis and Muscadinia* subgenera split ∼34 million years ago (Mya; 95% HPD: 32.77-36.31 Mya), similar to the age estimated from a smaller sample of complete genomes (Cochetel et al. 2023). The subgenus *Vitis* radiation dates to 22.3 million years ago (Mya; 95% HPD: 20.72-23.58 Mya), an intermediate value to previous estimates of ∼35 Mya (Nie et al. 2023) and ∼18 Mya (Cochetel et al. 2023). Second, we estimated the age of the Asian clade to be 17.7 Mya (95% HPD: 16.48-18.22 Mya, **Fig. 2A**), but our results were inconsistent as to whether the Asian species form a monophyletic clade within North American *Vitis*. In all analyses, *V. californica* was sister to the Asian clade, confirming it is the closest North American relative to Asian species (Talavera et al. 2023). Third, as expected, domesticated grapevine and/or its wild relative (ssp. *sylvestris*) formed a separate clade within the Asian group (**Fig. 1**; **Fig. 2A**). The domesticate was monophyletic, however (**Fig. 1)**, providing no clear evidence for two distinct domestication events (Dong et al. 2023) with these data despite the inclusion of *sylvestris* samples from throughout its geographic range.

Finally, the phylogenies supported three major *Vitis* clades that corresponded broadly to geographic distributions and ancestry inferred at K = 3 (**Fig. 1B**; **Fig. 2A**): the monophyletic Asian clade, a North American “Western/Central” clade, and a North American “Eastern” clade. The North American “Western/Central” clade contained *V. arizonica* and *V. girdiana,* which are endemic to the Southwest United States (US); *V. rupestris*, which is found primarily in the Ozarks but also in Oklahoma and Texas; and *V. riparia*, which is broadly distributed across the Central and Eastern US (Heinitz et al. 2019). The Eastern clade included *V. vulpina*, *V. labrusca*, and *V. aestivalis,* which are found across the Eastern US; *V. berlandieri* from Texas, New Mexico, and Arkansas; and *V. cinerea* from Florida and Texas (Heinitz et al. 2019).

### Rampant introgression correlates with geography

To formalize the complex admixture patterns (**Fig. 1**), we applied population genetic tests for introgression. We limited these analyses to the 19 species for which we had *n* > 3 individuals (**Table S4**), including three species of Asian origin. For each possible pair of species, we computed Patterson’s D-statistic and the related f4-ratio for all possible phylogenetically appropriate species trios (**Fig. 2A**) using *Muscadinia rotundifolia* as the outgroup (**Fig. 2B**). Overall, we tested 144 species pairs for asymmetric allele sharing based on 970 tests (**Table S5**). Of these, 122 of 144 (84.7%) species pairs yielded a maximum D value with a Z-score > 4 (FDR < 0.0001), indicating rampant historical introgression (**Fig. 2B**). Unsurprisingly, the three Eurasian *Vitis* lineages (*V. vinifera*, *V. sylvestris*, and *V. amurensis*) had weak introgression signals with the North American taxa, but introgression was pervasive among North American species pairs, many of which had evidence for introgression with one or more other species. For example, *V. aestivalis* had asymmetric allele sharing with three species in a separate clade (*V. riparia*, *V. rupestris* and *V. acerfolia*), perhaps reflecting gene flow prior to the divergence of the three species in this complex. The f4-ratio results were similar but provided an estimate of the admixture proportion for each species (**Fig. S8**). Across 19 species, the average f4-ratio was 0.14 (range: 0.01 to 0.78), meaning that on average roughly 14% of the genomes across our sample can be attributed to introgression. Consistent with their putative hybrid origins, *V. x doaniana* and *V. x champinii* had the highest f4-ratios; when they were removed from the dataset, the average f4-ratio was 0.12.

D-statistics are correlated across species pairs due to shared phylogenetic histories, so we explored alternative approaches to explore the history of introgression. One was TreeMix, which also revealed a history of pervasive introgression, including historical introgression events between *V. aestivalis* and the *V. acerifolia/riparia/rupestris* clade and also between the *V. arizonica/girdiana* clade and the *V. acerifolia/riparia* clade (**Fig. S9**). We also applied the f-branch (fb) statistic, which assigns excess allele sharing to specific branches on a species phylogeny. This analysis identified 42 branches with significantly elevated fb signals (fb > 0.1; FDR < 0.0001). Inferred introgression events included *V. arizonica* with *V. monticola*, *V. labrusca* with *V. vulpina*, and *V. californica* with both *V. arizonica* and *V. girdiana*. The strongest fb signals again confirmed the hybrid status of *V. x doaniana* and *V. x champinii* (**Fig. S10**).

Our study integrates across many species, making it possible to assess patterns that would be more challenging with just a few species. For example, it seems reasonable to hypothesize that geographically closely related species introgress more often. To test this idea, we used species distribution models (SDMs) to estimate the geographic range of 14 North American species, excluding the putative hybrid species (*V. x doaniana* and *V. x champinii*) to avoid confounding species-level patterns (**Fig. S11**). We first identified which species pairs have overlapping geographic distributions and plotted the distribution of D statistics for overlapping species compared to non-overlapping species. Overlapping species pairs had higher average D statistics (ANOVA: *F*_2,102_ = 36.48, *P*=1.12e-12; **Fig. 2C**). Second, we calculated the geographic centroid of each SDM and then, for each species pair, calculated the geographic distance (*d*_centroid_) between centroids. The pairwise D and *d*_centroid_ matrices were correlated, based on both multiple regression matrix analysis (P<0.001) and linear regression analysis (Pearson’s *r* = -0.41; *P* =2.35e-5; **Fig. 2D**).

We also explored the hypothesis that introgression facilitates adaptation to new ecological niches (Marques et al. 2019). We predicted that introgression should be more evident at geographic edges, where habitat tends to be more marginal and where the opportunity for hybridization may be more frequent. To test this idea, we used the inferred admixture proportions of each individual (**Fig. 1**), the sampling location of each individual when available (**Table S4**), and the location of each individual relative to the geographic centroid of its assigned species. After combining data across 233 individuals with location information that represent eight species, we found a significant positive correlation (Pearson’s *r* = 0.49; *P* = 2.12e-15) between the admixture proportion of individuals (as measured by 1-*Q*, where *Q* is an individual’s maximum ancestry proportion based on the admixture analyses; **Fig. 1**) and their normalized distances from geographic centroids. The strength of the relationship is diluted by the inclusion of species with little evidence for introgression; analyses on a per species basis yielded higher correlations for several species (e.g., *V. arizonica and V. rupestris*), with one as high as *R* = 0.70 **(Fig. S12)**. Altogether, these results show that *Vitis* has been profoundly shaped by widespread introgression, that introgression between species scales with geographic distance, and that these patterns are consistent with more admixture at geographic edges.

### Are Vitis x doaniana and Vitis x champinii long-term, stable hybrid species?

Our analyses consistently uncovered strong evidence of admixture within *V. x doaniana* and *V. x champinii*, reflecting their well-accepted hybrid origins (**Figs. 1 & 2B**). But is there evidence that they are distinct, historically stable species? We addressed this question by performing a series of population genomic analyses on hybrids and their parental donors. For simplicity, we present results based on *V. x doaniana* here in the main text, with parallel analyses reported in the supplement for the two distinct *V. x champinii* groups and a hybrid group of *V. arizonica* / *V. girdiana* individuals (**Fig. S14**).

We began by performing ADMIXTURE (K=2) analyses with *V. x doaniana* and its two parental species (*V. mustangensis* and *V. acerifolia*) (**Fig. 3A**), by measuring genome-wide heterozygosity for the parent-offspring trio (**Fig. 3B**) and by applying principal component analysis (PCA) to the genomic data (**Fig. 3C**). To aid the interpretation of the PCA and additional analyses, we included artificial ‘controls’ - i.e., *in silico* F1 hybrids produced by combining SNP variants from parental individuals (see Methods). The results showed that heterozygosity of *V. x doaniana* was higher than either parent, suggesting the possibility of recent hybridization; further, both PCA and admixture had *V. x doaniana* individuals as intermediate between donor species. We used triangular plots – which visualize ancestry proportions among three taxa (Wiens et al. 2025) – to evaluate these signals more precisely. As expected, the *in silico* F1 hybrids were placed at the apex of the plot, while the *V. x doaniana* individuals were largely aligned with the *in silico* hybrids or with regions consistent with early-generation backcrosses (**Fig. 3D**). The shallow age of these hybrids was further supported by DATES analyses, which estimates the timing of hybridization events for a single individual based on the size of recombination blocks (**Fig. 3E**). DATES estimated that the hybridization age of the 20 *V. x doaniana* individuals ranged from 1.58 ± 0.73 to 16.56 ± 4.83 generations, with an average of 2.95 ± 0.89 generations (**Fig. S13**). Chromosome painting for each individual revealed large, unbroken chromosomal segments from each parental species, a pattern consistent with recent hybridization and limited recombination (**Fig. 3F**). Finally, we considered whether geography could make *V. × doaniana* distinct from its parents, but the sampled *V. × doaniana* individuals were localized within the *V. acerifolia* SDM (**Fig. 3G**). Altogether, these data suggest that *V. x doaniana* is a recent hybrid. We uncovered qualitatively identical results for other hybrid groups (**Fig. S14**).

**Figure 3.**
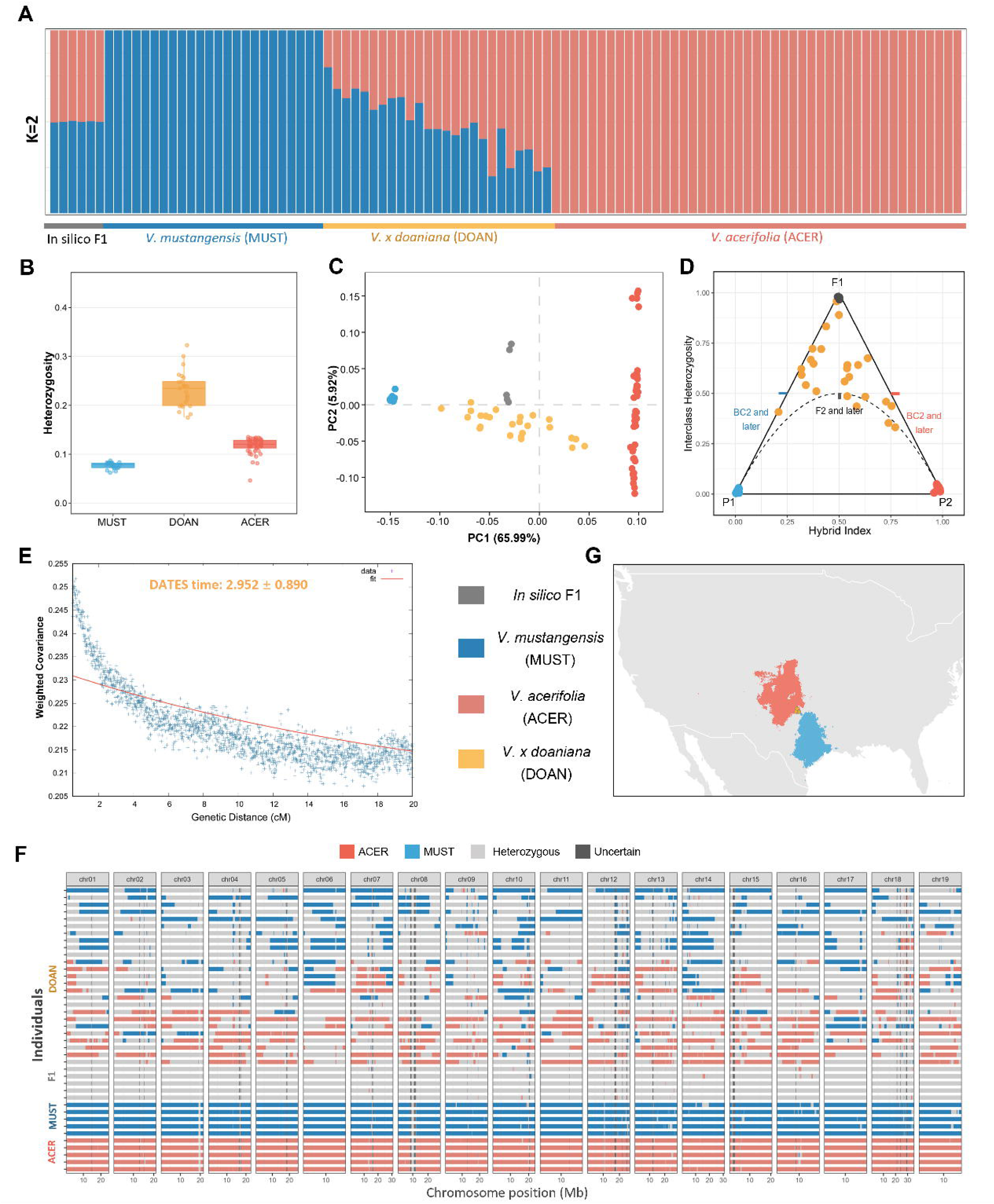
Genomic evidence for the recent hybrid origin of *Vitis x doaniana*. **(A)** Model-based admixture analysis (K=2) showing global ancestry proportions for *V. x doaniana* individuals, individuals from the hypothesized parents (*V. mustangensis* and *V. acerfolia*), and *in silico* F1 individuals. **(B)** Comparison of genome-wide heterozygosity of individuals. **(C)** Principal Component Analysis (PCA) of whole-genome SNPs of four groups. **(D)** Triangular plot of hybrid index (*S*) versus interclass heterozygosity (*H*). The plot compares natural *V. x doaniana* samples against diagnostic markers from parental species and *in silico* F1 controls, classifying individuals as early-generation hybrids (F1/F2) or backcrosses. The points above the dotted curve represent individuals that re estimated to be backcross 2 (BC2) individuals or more recent. **(E)** Inference of admixture generation timing using DATES. The curve displays the decay of weighted ancestry covariance against genetic distance (cM) for *V. x doaniana* populations. The average inferred age is 2.9 generations. **(F)** Chromosome painting visualization using WinPCA results of ancestry assignment. Each row represents the 19 chromosomes of a single individual. Genomic windows are colored by ancestry state: homozygous for *V. mustangensis* (blue), homozygous for *V. acerifolia* (red), or heterozygous (grey). (**G)** Geographic distribution showing complete overlap between the location of sampled *V. x doaniana* individuals and the species niche of one parent (ACER).

### Extensive repeated adaptation between *Vitis* species

Our multi-species sampling provides a unique opportunity to assess the repeatability of adaptive evolution, specifically whether adaptive evolution has targeted the same genomic regions across species. To explore these ideas, we focused on eight North American species with *n* ≥ 15 samples *(V. mustangensis, V. cinerea, V. berlandieri, V. girdiana, V. arizonica, V. rupestris, V. riparia and V. acerifolia)*, reflecting the fact that higher sample sizes provide more accurate inferences of selective sweep positions (Pavlidis et al. 2013). We scanned each species for signatures of recent positive selection using the Composite Likelihood Ratio (CLR) statistic. To differentiate adaptive signals from demographic noise, we established significance thresholds for each species using neutral coalescent simulations that incorporated species-specific demographic histories (**Fig. 4A, Fig. S15, S16**; see Methods). After we identified putative selective sweeps in each species, we defined a selective sweep as “shared” between species if the genomic intervals overlapped by > 60% of the sweep length.

**Figure 4.**
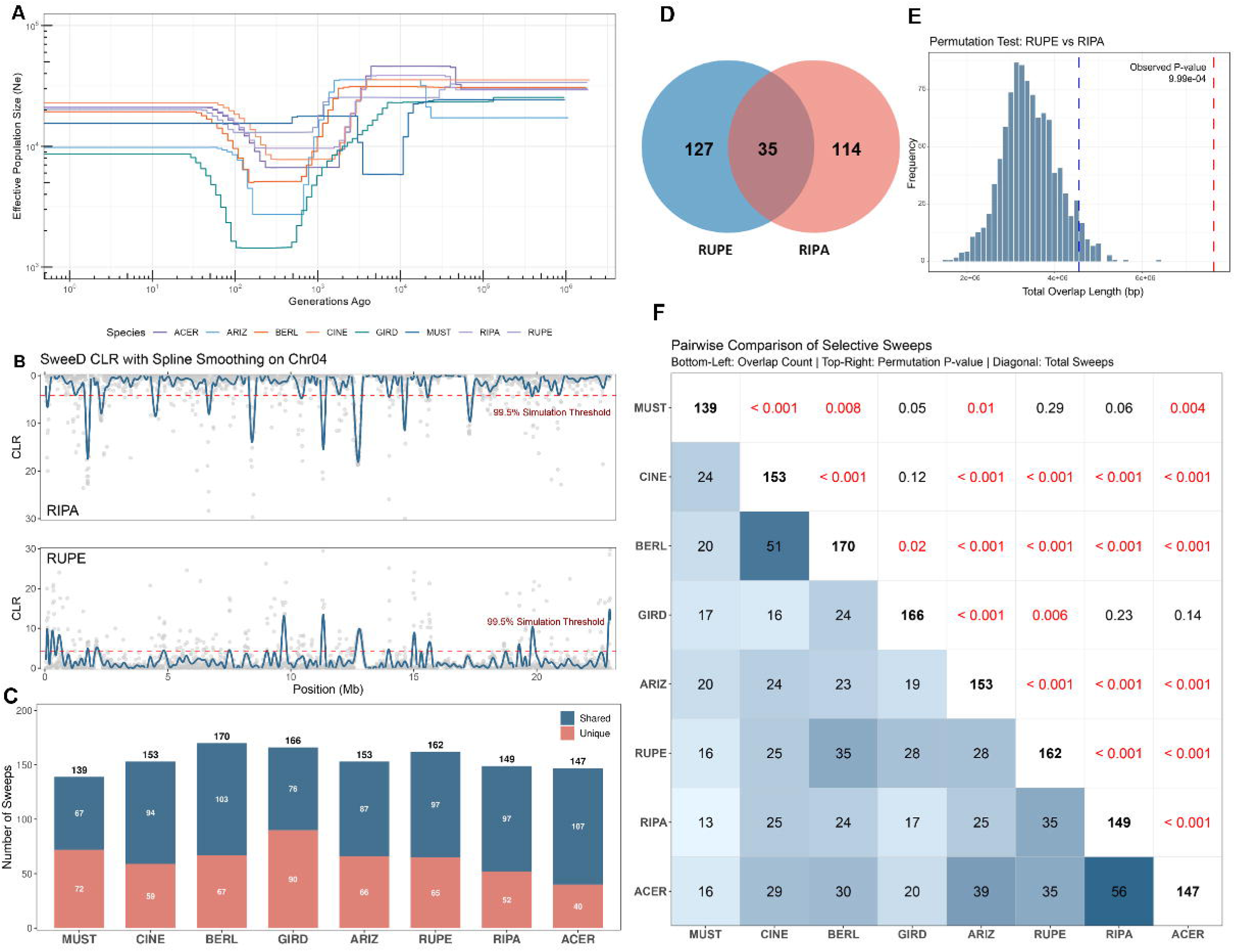
Identification and pairwise comparison of convergent selective sweeps among eight North American *Vitis* species. **(A)** Demographic history trajectories inferred for selected *Vitis* species, using mushi. The plot displays changes in effective population size over time (generations ago). **(B)** Examples of genome-wide scan for selective sweeps in species pair *V. riparia* (RIPA) and *V. rupestris* (RUPE) on chromosome 4. The solid blue line represents the spline-smoothed CLR profile used to identify peaks. The horizontal red dashed line marks the significance threshold, defined as the 99.5th percentile of neutral coalescent simulations. **(C)** Counts of unique and shared selective sweeps identified for each species. **(D)** Venn diagram showing the number of unique and shared selective sweeps between example species pair *V. rupestris* (RUPE) and *V. riparia* (RIPA). **(E)** Permutation test assessing the significance of putative sweep overlaps between RUPE and RIPA. The histogram shows the null distribution of overlap lengths expected by chance. The vertical red line indicates the observed overlap (*P* < 0.001); the vertical blue line represents *P* < 0.05. **(F)** Pairwise summary matrix of selective sweeps across eight *Vitis* species. Diagonal (bold): Total number of significant selective sweeps identified for each species. Lower Triangle (blue heatmap): The count of shared sweep regions between each species pair. Upper Triangle: P-values from permutation tests assessing whether the observed sharing is significantly greater than random expectation. Significant FDR-adjusted P-values (< 0.05) are highlighted in red.

We identified numerous regions consistent with positive selection (**Fig. 4B**). The total number of significant selective sweeps varied from 139 in *V. mustangensis* to 170 in *V. berlandieri*, representing between 5.06% and 7.13% of the total genome. The average sweep length was 174.5 kb, with 95% of sweeps between 23.5 kb to 505.7 kb (**Fig. 4F, Fig. S17, Table S6**). Each species contained unique (i.e., non-shared) sweeps, ranging from 40 in *V. acerifolia* to 90 in *V. girdiana*, but for six of eight species the majority of sweeps overlapped with those from other species (**Fig. 4C**). Examining the total of 734 shared sweeps, most overlapped between two (340) or three (213) taxa, and only one was shared among seven or more species (**Fig. S18**).

To test whether sweeps were shared more often than expected by chance between species, we performed permutation tests. The tests shuffled the locations of sweeps randomly across genomes while preserving their size. We illustrate the approach with *V. rupestris and V. riparia*, which shared 35 sweeps (**Fig. 4D**). After generating null distributions from shuffled sweeps, the observed length of overlapping sweeps was far greater than expected by chance (*P* < 0.001; **Fig. 4E, Fig. S19**). We repeated this analysis across all species pairs, revealing a pattern of repeated adaptation, with 22 of 28 pairwise comparisons significant (FDR-adjusted *P* < 0.05) (**Fig. 4F**). Not surprisingly, the strongest signals of repeatability were observed between phylogenetically close or geographically overlapping lineages - e.g., the *V. cinerea–V. berlandieri* pair and the *V. riparia–V. acerifolia* pair. Similarly, many of the non-significant comparisons included more distantly related or largely allopatric species, such as the *V.girdiana*-*V. riparia* pair. Finally, we examined gene content in sweeps, finding that a small proportion (2.66%) contained no genes. Combining genes from putative sweeps across all eight species, GO enrichment analysis identified pathways related to environmental responses and stress resistance (**Fig. S20**).

### Introgression is the dominant mode of repeated adaptation

The significant excess of shared selective sweeps among *Vitis* species raises at least two questions: does repeatability arise from independent *de novo* mutations, selection on ancestral standing variation, or adaptive introgression? And how do these mechanisms vary as a function of divergence time?

To distinguish among the three mechanisms, we applied a coalescent-based composite likelihood approach that models the probability of shared sweep signatures under the three distinct evolutionary mechanisms (Lee and Coop 2017). For each shared sweep region identified in our *Vitis* species pairwise comparisons, we computed the likelihood of *i*) neutrality, *ii*) independent *de novo* mutation, *iii*) migration (i.e., adaptive introgression), and *iv*) selection from ancestral, standing variation. We retained only those regions where the best-fitting adaptive model significantly outperformed the neutral model (composite likelihood scores difference, ΔCLE > 2), reducing the total number of shared sweeps to 663 from 734 (**Fig. 5A**). The analysis of these 663 shared sweeps suggested that convergent adaptation is rarely fueled by independent mutation; across all species pairs, only ∼5% (34 of 663) of shared sweeps were best explained by the model of independent *de novo* mutations (**Fig. 5A**). In contrast, 56% and 39% of shared sweeps better fit the adaptive introgression and standing variation models (**Fig. 5A-B**). It is worth emphasizing that these results were not due to the contribution of just a few species pairs, because migration was the numerically dominant model in 18 of 28 pairwise species comparisons (**Fig. 5C**).

**Figure 5.**
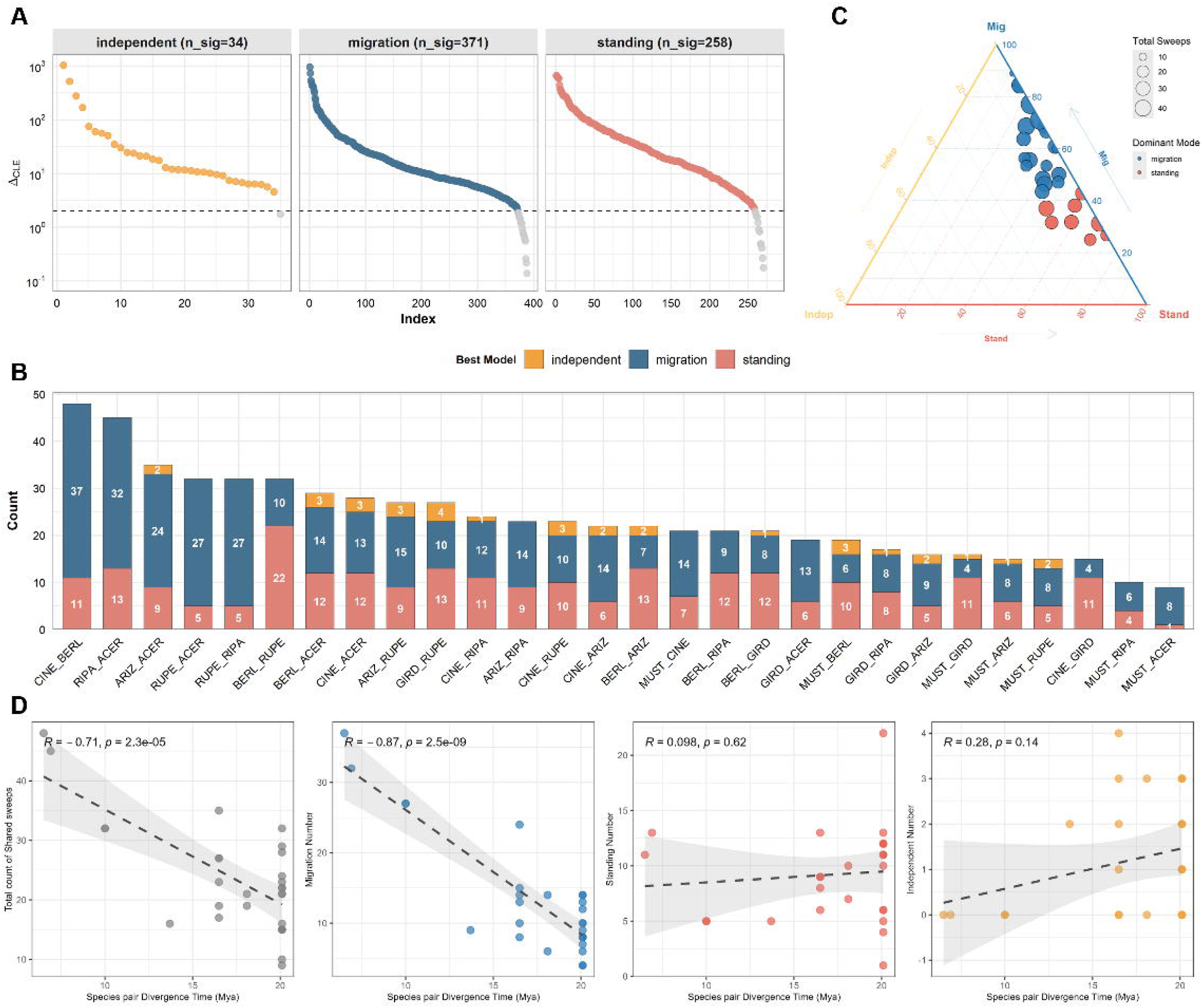
Inference of evolutionary mechanisms driving convergent adaptation in *Vitis*. **(A)** Assessment of model support for three modes of convergence: independent mutation, migration, and standing variation (i.e. ancestral sorting). The plots display the difference in composite log-likelihood between the best-fitting convergent model and the neutral model for each shared sweep region. The dashed horizontal line marks the significance threshold (ΔCLE > 2); only regions exceeding this threshold were considered to have significant support and were retained for downstream analysis. **(B)** Distribution of the best-supported evolutionary models across species pairs. The counts of shared sweeps best explained by migration (blue) and standing variation (red) far exceed those explained by independent *de novo* mutation (orange). **(C)** Ternary plot summarizing the proportion of shared sweeps assigned to each evolutionary mode for each species pair. Mig = migration model; Indep = *de novo* mutation model; Stand = standing variation model. **(D)** Correlations between divergence time (x-axis) and the convergence mechanism based on (from left to right) the number of total shared sweeps, migration sweeps, standing-variation sweeps, and independent-mutation sweeps.

We investigated whether repeated evolution scales with divergence time *(t*) (Bohutínská and Peichel 2024) by plotting the number of shared sweeps for each species pair against our *t* estimates (**Fig. 2A, Fig. S7**). The resulting plot revealed strong negative correlation (*R* = -0.71; *P* = 2.3e-5; **Fig. 5D**), and significance was confirmed by Phylogenetic Generalized Linear Mixed Model (PGLMM) analysis (*P* = 4.4e-4). We also plotted each of the three categories separately. There was no statistically supported relationship between *t* and either the number of *de novo* sweeps or the number of sweeps based on ancestral variation. In contrast, the number of adaptive introgression events declined over time (linear regression, *R* = -0.87, *P* = 2.5 x 10^-9^; PGLMM, *P* = 4.0e-5) but did not drop to zero because even species that diverged at the origin of the subgenus 22 ∼Mya shared overlapping sweeps (**Fig. 5D**).

## DISCUSSION

Many questions in evolutionary biology require accurate characterization of the patterns and rates of species diversification. Population genomic datasets spanning multiple species are particularly valuable, because they can simultaneously inform about phylogenetic relationships, divergence times, the extent of hybridization, the genomic basis of trait evolution and even landscape-scale evolutionary patterns (Bock et al. 2023; Combrink et al. 2025). Datasets of this type are likely to be particularly informative in plants, where hybridization has long been proposed as a driver of adaptive radiation (Anderson and Stebbins 1954) and affects species relationships. Despite their importance, few systems currently offer whole-genome resequencing resources that include population sampling across multiple species. This *Vitis* dataset contains, to our knowledge, the most wild plant species with explicit population genomic sampling from any plant genus or family to date.

### Introgression and reticulate species histories

Our analyses have an overarching conclusion: the evolutionary history of North American *Vitis* has been shaped dramatically by hybridization and introgression. The prevalence of introgression was evident from signals of asymmetric allele sharing across the genus, based on multiple metrics (**Fig. 2B**). For example, Patterson’s D-statistic identified significant asymmetric allele sharing in 84.7% of 144 species pairs, and the f4-ratio metric estimated admixture proportions averaging 0.14 across 19 species samples. TreeMix similarly supported extensive historical gene flow, while f-branch analyses localized excess allele sharing to dozens of branches on the species phylogeny (**Fig. 2A, Figs. S8–10**). These results extend previous work in *Vitis* (Zecca et al. 2019; Morales-Cruz et al. 2021; Nie et al. 2023; Xiao et al. 2025) and drive home that introgression has not only been common but pervasive.

The pattern of introgression is not random, however, because it is structured by geography and ecology, as measured by bioclimatic variables integrated into SDMs. For example, we found that the magnitude of D statistics between species pairs was elevated for species that overlapped in geographic niche space and also negatively correlated with geographic distances (*d*_centroid_) between species (**Fig. 2C, 2D, Fig. S11**). Relationships between introgression metrics and geography have not been investigated widely, but surveys of suboscine birds and wild tomatoes have also found introgression signals to be higher for species in close geographic proximity (Hamlin et al. 2020; Singhal et al. 2021). It is difficult to conclude that geography is causal, since introgression is likely to be confounded with other factors like phylogenetic relationships, but it is encouraging that geography is related to genome wide introgression statistics.

Similar geographic considerations apply among individuals within species, because *Vitis* individuals of admixed ancestry are more likely to be located at species’ margins (**Fig. 2E & S12**). This result is consistent with introgression facilitating persistence in marginal habitats at the edge of species’ distribution (Pfennig et al. 2016). Similar phenomena have been documented in *Quercus* (oaks), where introgression promotes adaptation and establishment in edge populations (Leroy et al. 2020; Parker et al. 2025); in *Populus* (poplars), where introgression has facilitated adaptation to high latitudes (Rendon-Anaya et al. 2021); and in *Carex* species, where hybrid individuals occupy unique environmental conditions (Hodel et al. 2022). It is worth noting, however, that effects at the edge of species’ distributions are likely to be nuanced. For example, a landscape-scale study of *V. arizonica* found that populations at the edge of range expansion have more deleterious alleles and fewer adaptive alleles than geographically central populations, likely due to population histories of dispersion and associated effects like small population sizes (Fiscus et al. 2025). Our point is that introgression may aid persistence and adaptation in marginal habitats, but it may be insufficient to erase genome-wide signatures associated with range expansion and marginal habitat.

One salient question is whether *Vitis* represents a typical plant adaptive radiation with respect to the prevalence of introgression. Some biological features may make it atypical. One is the interfertility of species. All *Vitis* species are facultatively outcrossing, which promotes cross-breeding, and are typically pollinated by generalist insects, which may again facilitate cross-species hybridization. They are also long-lived perennials, which can allow the overcoming of any leaky temporal mating barriers when environmental conditions fluctuate from year to year (Gaut et al. 2015; Farnitano et al. 2025). The presence of extensive zones of sympatry is also likely to elevate introgression effects relative to systems with more geographic isolation. In *Vitis,* even species from distinct phylogenetic clades – such as the “Central/Western” clade and the “Eastern” clade – have spatial overlap (**Fig. S11**), creating a vast landscape of potential contact zones. Finally, studies of complete genomes throughout the subgenus have discovered few of the genomic features – such as prominent inversions or extensive variability in genome size – that might impede hybridization on a genomic level (Cochetel et al. 2023). Altogether, these considerations suggest *Vitis* could be atypical, but similar data from more systems are needed to make this argument with any certainty.

The interconnected nature of *Vitis* species complicates phylogenetic inference, but the economic and ecological importance of the genus has nevertheless made it a frequent target for phylogenetic analysis (Jaillon et al. 2007; Goremykin et al. 2009; Wan et al. 2013; Liu et al. 2016; Wen et al. 2018; Talavera et al. 2023). These analyses have generally failed to reach a consensus about species’ relationships (Miller et al. 2013), probably for reasons that now seem obvious: phylogenetic discordance between plastid and nuclear trees (Larson et al. 2026), along with limited population sampling in most previous studies. Population sampling is a potential issue because if only one or two individuals are studied per species, those individuals may reflect admixture events more than species’ divergence. Of course, there is no simple solution to the problem of assessing species relationships when there is reticulate evolution. Here we chose to generate a total evidence tree based on all of the available data (**Fig. 1**) and also constructed species-level trees after filtering notably admixed individuals (**Figs 2A & S4-6**). The major inferences largely mirrored those from another recent study based on 43 species that identified 9 major clades (Talavera et al. 2023). Species sampling was not identical between the two studies, but our RaXML, SVDQuartets, and reference-free Kmer trees recapitulated their nine main groups as well as most of the relationships among groups, with the prominent exception of lingering uncertainty about the origin of the Asian clade (which varied among our trees and also among previous phylogenetic studies). There is bound to be phylogenetic uncertainty in any group with reticulation, but this and previous work (Talavera et al. 2023) are beginning to yield a consensus for *Vitis*.

### Scant evidence that hybridization leads to speciation

Introgression is common in *Vitis*, but is there any evidence that hybridization has created distinct species? Theoretically, for a hybrid lineage to be distinct and stable, it needs to evolve some degree of reproductive isolation from its parents or occupy a distinct ecological niche that promotes persistence (Schumer et al. 2014; Long and Rieseberg 2025). Genomically, ancient homoploid hybrid species are expected to show a fine-scale mosaic of parental ancestry produced by recombination over many generations, thereby forming a genetic cluster distinct from recent-generation hybrids (Wang et al. 2025; Wiens and Colella 2025).

*Vitis* contains two taxa, *V. x doaniana* and *V. x champinii*, that have been recognized as hybrids (Heinitz et al. 2019; Zecca et al. 2019), but their genomic histories have not been evaluated. One notable finding is that *V. x champinii* is not genetically uniform (**Fig. 1**), because there are two groups with apparently different admixture donors. This observation explains the disagreement as to whether *V. rupestris* or *V. monticola* is one of the hybrid parents (Heinitz et al. 2019), because both are true. Nonetheless, genomic evidence supports that all of the studied hybrid taxa are the product of recent (and perhaps ongoing) hybridization. For example, ancestry-based approaches placed hybrid individuals as similar to *in silico* F1 controls, triangular plots were consistent with early-generation backcrosses, and age estimates were recent (i.e., from 2 to 17 generations, representing roughly < 10 to 60 years) (**Fig. 3 & S13**). Chromosome painting also revealed large, unbroken ancestry tracts from each parental species, which is again consistent with recent origins and limited recombination since admixture. No genomic regions obviously differentiated all hybrid individuals from their parents (**Fig. 3F**), further suggesting the hybrids have not differentiated as a lineage.

These results contrast markedly with genomic patterns from other homoploid hybrid species. For example, species in the genus *Ostryopsis* are estimated to have originated ∼1.8 Mya and are readily distinguished by parental chromosomal blocks that have been reassorted into characteristic mosaic patterns (Z. Wang et al. 2021). Similar evidence of ancient stabilization and genomic reorganization has been found in hybrid lineages of sunflowers (Owens et al. 2023), chestnut trees (Sun et al. 2020), bears (Zou et al. 2022) and monkeys (Wu et al. 2023). Even in a relatively young hybrid species like *Senecio squalidus*, which is ∼300 years old, the genomes display evidence of fragmented ancestry and reduced heterozygosity (Nevado et al. 2024).

Altogether, our genome-wide analyses show that neither *V. x doaniana* nor *V. x champinii* are genetically persistent (i.e., ancient) taxa. However, either could be a nascent hybrid species if they are either reproductively isolated or ecologically distinct from their parents. Although we did not perform controlled crosses, the common use of both taxa in grapevine genetic analyses and breeding [e.g., (Zou et al. 2023)] suggests there is little, if any, reproductive isolation. As for ecological distinctness, the sampled *V. x doaniana* individuals fell within the SDM of *V. acerifolia* (**Fig. 3G**), suggesting *V. x doaniana* is not ecologically distinct at this scale. Altogether, we have no evidence to suggest that any of the four *Vitis* hybrid groups are distinct or even nascent species. We believe, however, that our analyses offer an important negative result: hybrids in nature may not present genomic evidence for homoploid hybrid speciation events, even when they are commonly recognized as distinct hybrid taxa.

That is not to imply, however, that *V. x doaniana* and *V. x champinii* are inconsequential for *Vitis* evolution and diversification. They represent examples of hybrid swarms that are common in *Vitis*, given that we detected a hybrid group in *V. x doaniana,* two groups of *V. x champinii* and at least one other group between *V. arizonica* and *V. girdiana* (**Fig. 1**). These hybrids can be agronomically valuable; for example, *V. x champinii* is commonly used for rootstock development, in part because it has the “high vigour”(Chen et al. 2024) typical of heterosis. In the larger context of an adaptive radiation, hybrid swarms can reinforce species boundaries (Groh and Coop 2024) but they are also hypothesized to contribute to the production of novel genetic combinations that help drive rapid diversification (Grant and Grant 2019; Marques et al. 2019; Combrink et al. 2025). If the latter is true, these swarms contribute to a perspective that portrays North American *Vitis* as a series of interconnected lineages - i.e., a ‘syngameon’ (Seehausen 2004). Moreover, we posit that the hybrid groups represent a critical feature of an ongoing adaptive radiation(Combrink et al. 2025) in *Vitis* and may be more common in other plant adaptive radiations than currently recognized.

### Repeated adaption is fueled by introgression

Our dataset has provided a unique opportunity to assess parallel evolution, or adaptive repeatability. Several recent studies have measured repeatability in plant systems, generally using one of two approaches. The first is to test for genotype-environmental associations (GEA) and assess whether homologous/orthologous genes exhibit associations across different lineages (Yeaman et al. 2016). For example, Whiting et al. (2024) performed GEA across 25 plant species representing divergence times from 2.5 Mya (Nocchi et al. 2024) to > 300 Mya. They found that genes with specific functions were over-represented, suggesting that some core physiological pathways have been commonly targeted by natural selection throughout plant evolution (Whiting et al. 2024). The second approach, which we have used here, is to compare regions of putative selective sweeps among species. The advantages of this approach include an explicit genomic context and the ability to detect phenomena outside of coding regions. Relevant examples for this approach come from studies in *Zea*, where there is evidence of parallel genetic adaptation to high elevation across subspecies and populations (L. Wang et al. 2021; Tittes, Lorant, McGinty, et al. 2025). Note, however, that the genus *Zea* radiated only ∼150,000 years ago (Ross-Ibarra et al. 2009; Chen et al. 2022), representing at best ‘a blink of an eye’ on an evolutionary timescale.

*Vitis* is a suitable model for studying repeatability because it is speciose, has highly collinear genomes across species (Cochetel et al. 2023), and represents an appreciable expanse of evolutionary time (**Fig. 2A & S7**). Our analysis of selective sweeps across North American *Vitis* extends our previous research (Morales-Cruz et al. 2021) which was based on fewer species. Here we have increased the number of species, improved the specificity of selective sweep mapping by explicitly incorporating demographic inference, and focused explicitly on potential mechanisms of repeatability. Most species pairs (22 of 28 = 78%) shared significantly more selective sweeps than expected by chance (**Fig. 4**), suggesting extensive parallel evolution. The interpretation of 78% must be tempered by understanding that the statistical power to detect sweeps varies among species, that there is statistical non-independence across species pairs, and that our genome-wide permutations could be anti-conservative if sweeps are limited to just a few gene-rich or recombination-poor genomic regions. We believe it unlikely, however, that genome architecture alone accounts for the striking degree of parallelism. One reason is that ∼35% of the reference grapevine genome is gene rich (Shi et al. 2023), meaning the proportion of genome encompassed in sweeps is still small compared to the gene rich region. Furthermore, if shared sweeps are caused by genomic architecture, one expects sweeps to be shared across most or all taxa, reflecting the highly conserved architecture across *Vitis* genomes (Cochetel et al. 2023). However, only one of 743 putative sweeps overlapped across > 6 species, while 75% overlapped between only two or three species (**Fig. S18**).

The high degree of shared sweeps raises the question of mechanism: does repeatability arise from independent *de novo* mutations or from shared pools of genetic variation? A coalescent-based modeling framework (DMC) (Lee and Coop 2017; Tittes 2020) has documented overwhelming support for models of shared ancestry (**Fig. 5**). Specifically, we identified 629 shared adaptive regions that were better explained by the two shared ancestry models – adaptive migration or ancestral sorting – compared to only 34 attributable to independent mutations (**Fig. 5A**). Among shared ancestry models, our analyses estimate that repeatability occurs ∼1.45 more often via adaptive introgression (i.e., migration) than through ancestral sorting. This general pattern aligns with *Zea*, where selective sweeps were shared among populations most frequently via migration (Tittes, Lorant, Mcginty, et al. 2025). It must be remembered, however, that our sweeps were categorized by the model with the most support, as opposed to statistically significant differences in model likelihoods. For example, among the 663 sweeps that favored the non-neutral models, 140 did not show statistically significant support (i.e., ΔCLE < 2) for the best-fitting model over the second-best alternative. There is thus some uncertainty in model categorization. Nonetheless, the inescapable conclusion is that shared ancestry is the principal driver of adaptive repeatability in *Vitis*, and the evidence points to adaptive introgression as the most common source for parallel events.

One expects the drivers of shared ancestry to dissipate over time (Bohutínská and Peichel 2024), because more distant lineages should share fewer segregating ancestral variants and/or have higher impediments to gene flow, due to either geographic or reproductive isolation. Thus far, however, the relationship between repeatability and divergence time has been tested in only a few systems (Bohutínská et al. 2021; Bohutínská and Peichel 2024). We could address this issue here, because the eight species that we used in DMC analyses ranged in divergence time (*t*) from 6.5 to 22 Mya (**Fig. S7**). Overall, there was a strong negative relationship between *t* and the number of shared sweeps, and the relationship was particularly pronounced for sweeps attributed to migration (**Fig. 5D**). These observations substantially expand our understanding of the pace and distribution of repeatable adaptive events.

Taken together, our results support that repeated adaptation in *Vitis* is shaped by a shared pool of variation that is maintained by both ongoing gene flow and through ancestral polymorphism. The dominance of shared ancestry in adaptation supports a “repeated sorting” framework, where similar phenotypes evolve in related lineages via the selection of identical by descent alleles (Waters and McCulloch 2021; Chaturvedi et al. 2025). The contribution of these mechanisms depends in part on phylogenetic (Waters and McCulloch 2021) and, as we have shown, geographic scales. In the diversification of North American *Vitis*, it is clear that “reinventing the wheel” from *de novo* mutation is evolutionarily inefficient compared to borrowing (via introgression) or inheriting (from standing variation).

## MATERIALS AND METHODS

### Genome sequencing and SNP calling

Our dataset consists of sequencing reads from published work and newly generated sequences. The previously published data are available from the NCBI Short Read archive at PRJNA393611(Liang et al. 2019b), PRJNA842753 (Morales-Cruz et al. 2023), PRJNA321480 (Magris et al. 2021), PRJNA984685 (Cochetel et al. 2023) and PRJCA009314(Dong et al. 2023). New Illumina whole genome sequencing reads from 114 accessions were generated following the methods described in(Morales-Cruz et al. 2021) and were deposited under PRJNA1424665. Detailed information about sources, species and accession sampling locations can be found in **Table S1**.

Sequencing reads were adapter and quality trimmed using Trimmomatic 0.39(Bolger et al. 2014) with the following options: “ILLUMINACLIP:“$ADAPTERSPE”:2:30:10 LEADING:3 TRAILING:3 SLIDINGWINDOW:4:20 MINLEN:60”. The resulting reads were then mapped to haplotype 1 of the *V. arizonica* v.2.0 genome assembly (Morales-Cruz et al. 2023) using bwa-mem 0.7.12-r1039(Li 2013) with the default options. Alignments were sorted and converted to binary format with samtools 1.10(Danecek et al. 2021). Duplicates were marked with picard MarkDuplicates integrated into GATK 4.2.6.1(McKenna et al. 2010; DePristo et al. 2011; Poplin et al. 2018). Samples were genotyped using GATK HaplotypeCaller in GVCF mode prior to joint calling with GenotypeGVCFs. Raw variants were filtered with bcftools 1.17(Danecek et al. 2021) to retain only biallelic SNPs with site quality of 20 or better, that passed the GATK recommended hard filters (exclude “QD < 2 | FS > 60 | SOR > 3 | MQ < 40 | MQRankSum < -12.5 | ReadPosRankSum < -8.0”), had a MAF >= 0.01, and had site depth greater than the mean plus standard deviation of depth across all sites. Finally, we filtered for missing data by removing individuals with greater than 70% missing calls across sites and sites with greater than 5% calls missing across individuals.

### LD pruning and admixture

SNPs were pruned for linkage disequilibrium using PLINK 2.0 (Purcell et al. 2007; Chang et al. 2015) with a pairwise sliding-window approach (50 SNP windows, 10 SNP step size) and an R^2^ threshold of 0.20. The pruned SNP set was then filtered to remove variants with a minor allele frequency (MAF) < 0.05. PCA was performed in PLINK 2.0 using a variance-standardized relationship matrix. Admixture analysis on genotype likelihoods was performed with NGSAdmix v32 (Skotte et al. 2013) considering K = 1 to 20 subpopulations at 20 replicates per K. Plink-formatted genotypes were converted to beagle format with ANGSD v0.940 (Korneliussen et al. 2014), which was used as input to NGSAdmix. The optimal number of ancestral populations was determined by calculating the delta K statistic (Evanno et al. 2005) (**Fig. S2**). Admixture data was then processed and visualized using the R package pophelper (Francis 2017).

### Phylogenetic inference

We constructed maximum likelihood (ML) phylogenetic trees using linkage disequilibrium (LD)-pruned SNP datasets from all samples. IQ-TREE v2.2.0 (Minh et al., 2020) was used to infer phylogenies under the best-fit nucleotide substitution model (GTR+R7), as identified by built-in ModelFinder (Kalyaanamoorthy et al. 2017). Bootstrap values for each node were calculated using the ultrafast bootstrap method [UFboot2; (Hoang et al. 2018)] with 1,000 replicates. The resulting trees were then visualized using the R package ggtree v4.0.4 (Yu 2020). For all trees we designated *Leea guineensis* as the outgroup.

For species-level phylogenetic inference, we first removed samples potentially influenced by admixture using two complementary approaches. First, we screened for outliers within each species using principal component analysis. For species represented by three or more samples, we calculated pairwise Euclidean distances among individuals in multidimensional PC space and computed modified Z-scores based on each sample’s median distance to others from the same species. Samples with a modified Z-score > 3.5 were flagged as putatively admixed. Second, we manually examined ancestry estimates from the admixture analysis to confirm PCA-identified outliers and to evaluate species represented by fewer than three samples. Samples flagged by either approach were considered putatively admixed and were not considered in subsequent analyses.

From the remaining samples, we randomly selected up to three individuals per species to represent intraspecific genetic diversity on the phylogeny. With this curated dataset, species relationships were verified using two approaches: a concatenated SNP tree inferred with RAxML-ng v1.1.0 (Kozlov et al. 2019) using the GTR+G model of molecular evolution with 100 bootstrap replicates and a coalescent-based inference using SVDquartets in Paup v4a166 (Swofford 2002) by evaluating all quartets and using 1000 bootstrap replicates. To estimate divergence times among species, we constructed a clock-calibrated phylogeny using BEAST2 v2.7.7 (Bouckaert et al. 2014). We selected the GTR nucleotide substitution model with four gamma-distributed rate categories and applied a relaxed molecular clock. A Calibrated Yule prior was specified for tree branching processes with a fixed topology from SVDquartets. Markov Chain Monte Carlo (MCMC) sampling was performed for 10 million generations, with tree states sampled every 1,000 generations and discarding the initial 10% as burn-in. Fossil calibration priors were defined using a log-normal distribution based on previously published divergence estimates between subgenera *Vitis* and *Muscadania* (Wan et al. 2013). The posterior distribution of calibrated trees was visualized using DensiTree v2.6.4 (Bouckaert 2010).

To generate a reference-free phylogeny, K-mer–based sketches were generated for each sample from the trimmed sequencing reads using mash v2.3 with default parameters (Ondov et al. 2016). Pairwise mash distances were computed using *mash dist*, yielding a distance matrix summarizing genome-wide sequence similarity. This matrix was used to infer a neighbor-joining tree using the *nj* function in the R package ape v1.3 (Paradis and Schliep 2019). The plastid genomes were assembled from trimmed sequencing reads using getOrganelle v.1.7.7.0 (Jin et al. 2020) with default parameters. For samples in which a complete plastid genome was not recovered, assemblies were reattempted using the parameters ‘-R 30 -w 95 –disentangle-time-limit 7200 –max-reads inf’, with the *Vitis vinifera* plastid genome (NCBI RefSeq NC_007957.1) used as a seed reference.

Resulting plastid assemblies were annotated with CHLOE (https://chloe.plastid.org/annotate.html) via its python API. Based on these annotations, coding sequences corresponding to the 101 plastid genes present in all 551 assemblies were extracted using bedtools v.2.31.1 getfasta (Quinlan and Hall 2010). Sequences for each gene were aligned independently using MAFFT v.7.505 (Katoh and Standley 2013) with the options ‘--maxiterate 2 –retree 2’. Individual gene alignments were concatenated into a supermatrix using seqkit (v2.4.0) concat (Shen et al. 2024) The concatenated alignments were then used to infer a maximum likelihood phylogeny using RAxML-ng v1.1.0 using the GTR+G model of molecular evolution and 100 bootstrap replicates. We calculated the Generalized Robinson-Foulds (GRF) distance between tree topologies using the R package TreeDist v.2.11.1 (Smith 2021). This metric quantifies tree discordance on a normalized scale from 0 (identical topologies) to 1 (no shared information).

### Detecting Introgression between Species

To investigate introgression between various *Vitis* species, we initially applied Patterson’s D statistic (Durand et al. 2011) with a four-taxon topology using *M. rotundifolia* as the outgroup (O). We tested whether any of two taxa (P1 or P2) shared an excess of alleles with a potential introgression partner (P3). D-statistics for all possible combinations of species trios within met inferred phylogenetic relationships were computed using the Dtrios module in Dsuite v0.5 (Malinsky et al. 2021) using default parameters. Statistical significance and standard errors for D-values were obtained using block jackknifing for non-overlapping contiguous genomic blocks. We also calculated the overall f4-admixture ratio, which quantifies the proportion of introgressed genome segments from a donor lineage, utilizing the admixr package v0.10 with default parameters (Petr et al. 2019). We also calculated the f-branch statistic from Dsuite fbranch module with default parameters (Malinsky et al. 2021), which assigns migration events to specific internal branches on the phylogeny. We applied Treemix v1.3 (Pickrell and Pritchard 2012) by constructing a phylogenetic tree without migration events using the LD-pruned SNP data and *M. rotundifolia* as the outgroup. We evaluated models by adding between 1 and 19 migration edges. The optimal number of migration events was determined using optM package v0.1.5 (Fitak 2021).

### Species Distribution Modeling

We generated species distribution models (SDMs) for 14 *Vitis* species from North America, emphasizing present-day distributions and expanding on statistical approaches previously employed to predict future SDMs (Zhang et al. 2025). Geographic occurrence data were obtained from the Global Biodiversity Information Facility (GBIF) (User 2025). Current bioclimatic variables corresponding to occurrence data were downloaded from WorldClim2 (Fick and Hijmans 2017), based on mean observations from 1970 to 2000. After obtaining occurrence and bioclimatic variables, we followed the methodology described in two previous publications (Aguirre-Liguori et al. 2021; Fiscus et al. 2025), with some modifications. Briefly, for each species we first cleaned the occurrences using the R package CoordinateCleaner v2.0 (Zizka et al. 2019), to remove potential erroneous locations. Next, we used the *M_simulationR* function in the package Grinnell package v.0.0.22 (Machado-Stredel et al. 2021), and the *mask* function in terra package (Hijmans et al. 2024) to delimit the accessible area (M Layer) for each species. The M layer was built based on simulated individual dispersal and the overlap between the remaining occurrence records and the world’s terrestrial ecoregions (Olson et al. 2001). Additionally, for the bioclimatic variables, we used the *CorSelect* function of the fuzzySim package (Barbosa 2015) to select uncorrelated variables (R < 0.8) based on a variance inflation factor.

The cleaned occurrences and the selected bioclimatic variables were used to build SDMs with the R package BIOMOD2 v. 4.2-4 (Thuiller et al. 2009). For each species, we ran 10 bootstrap replicates, using 70% and 30% of locations for training and testing with each of three algorithms (Random Forest, Generalized Additive Models and Maxent) to build 30 models per species. Finally, we generated ensemble models by weighting individual models according to their ROC (>0.8) and TSS (>0.6) values, producing final binarized and projected distributions. One species, *Vitis berlandieri,* had no entries in GBIF; we only were able to obtain records from the locations that were sampled for this study. We were therefore unable to run SDMs for this species but instead estimated its geographic distribution using a convex hull approach with the *hull* function in the terra v.1.8-70 package (Hijmans et al. 2024).

### Geographic centroids, species overlaps and landscape-scale admixture analyses

Once binarized SDMs were obtained, we estimated the geographic centroid of each species using the *centroid* function in the terra package (Hijmans et al. 2024). For each pixel in the species distribution *SpatRaster*, we extracted the geographic coordinates and calculated the Euclidean distance from each pixel to the species’ geographic centroid. The values were then rasterized to produce a *SpatRaster* representing the distance of each location to the geographic centroid. We exclude the putative hybrid species (*V. × doaniana* and *V. × champinii*) pairs to avoid confounding species-level patterns. We also analyzed the geographic overlap and distance between geographic centroids for a total of 105 possible pairs of species with SDMs available. For each species pair we identified the area where the species overlapped and used the *expanse* function in the terra package to obtain the overlap area in Km^2^. Species pairs were categorically defined as i) overlapping North American species (SDM overlap > 0 km²);ii) non-overlapping North American species; iii) species contrasts that included Eurasian taxa (*V. vinifera*, *V. sylvestris*, and *V. amurensis*). We also estimated the geographic distance between the centroids (*d*_centroid_) of each pair of species, using the *distGeo* function in the Geosphere v.1.5-18 package (Hijmans et al. 2019). We compared the maximum D and pairwise matrices using Multiple Regression Models, as implemented in the *MRM* function in the Ecodist package (Goslee 2026). Linear models were calculated in R.

To assess the relationship between admixture proportions of individuals and their geographic location, we calculated metrics that summarize the admixture proportion for each individual. We used two: 1) the maximum proportion from a single species (maxQ) based on population structure at K = 13; and 2) the effective number of contributing genomes [effN = exp(H), where H is the Shannon entropy estimated as 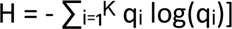. We report results based on maxQ, but results with effN were qualitatively identical. For the 233 individuals with available sampling locations (**Table S4**), we calculated the Euclidean distance to their geographic centroids using distGeo (Hijmans et al. 2019). We estimated the association between the admixture metrics and their distance to the geographic centroid using linear models as implemented with the lm function in R.

### Investigating the History of Putative Hybrid Species

To investigate the history of hybrid groups, we assumed the parentage of each group from admixture results and/or from the existing literature [e.g, (Heinitz et al. 2019)] (**Fig. 3 and Fig. S14**). For each of four hybrid groups (**Table S4**), we generated eight *in silico* F1 hybrids as a control group by combining SNP variants from randomly chosen six individuals of each assumed parental species. We performed the same analyses on each of the four sample groups. The first was PCA and admixture analyses, following the procedures described above. We also used the Hlest method implemented in triangulaR (Wiens et al. 2025), which explicitly assesses hybrid status by estimating parental individuals, backcrosses, F1, and later hybrid generations (≥F2) based on ancestry (S) and heterozygosity (H) at diagnostic loci (Wiens and Colella 2025). Ancestry-informative markers were selected by identifying loci with allele frequencies ≥0.9 in one parental species and ≤0.1 in the other.

The timing of hybridization events for each hybrid individual were estimated using DATES v753 (Distribution of Ancestry Tracts of Evolutionary Signals) (Chintalapati et al. 2022). DATES infers the time of population admixture by converting the average length of ancestry blocks into generations, based on a recombination rate of 6.4e-8 per base pair per generation (Zhu et al. 2018). The analysis was performed using the following parameters: binsize = 0.00005, maxdis = 1.0, runmode = 1, chithresh = 0, mincount = 1, qbin = 50, lovalfit = 0.1, and runfit, afffit, and zdipcorrmode set to YES. We inferred local ancestry across the genome for each hybrid-parent group using the window-based PCA, WinPCA v1.2 (Blumer et al. 2025), which performed PCA analyses within sliding genomic windows (200 Kb window 10 Kb step) across all 19 chromosomes. We used ggplot2 to generate chromosome painting diagrams.

### Demography, Selective sweep identification and population genetic simulations

We studied population genetic properties, including demography and selective sweeps, of eight *Vitis* species with n > 15 individuals **(Table S4)**. We used a three-tiered approach to detect selective sweeps. First, we inferred the demographic history of each species. We used mushi v0.2 (DeWitt et al. 2021) to estimate changes in effective population size (*Ne*) over time. The unfolded site frequency spectrum (SFS) was inferred for ten randomly selected individuals per species using est-sfs (Keightley and Jackson 2018), with ancestral states polarized using *M. rotundifolia* as an outgroup. To prevent overfitting with limited sample sizes, regularization parameters were set to ridge_penalty = 1e4, alpha_spline = 1e4, and trend_penalty = 1e2. Analyses assumed a mutation rate of 5.4e-9 per nucleotide per year (Liang et al. 2019a) and a generation time of three years (Zhou et al. 2017).

Second, we generated neutral replicates for each species using coalescent simulations msprime (Baumdicker et al. 2022)under species-specific demographic models. For each species, we set simulation parameters using effective population size trajectories inferred from mushi, incorporating ancestral population sizes, bottleneck population size and timing, and exponential growth where applicable. Simulations assumed a mutation rate of 5.4e-9 per nucleotide per year and a recombination rate of 6.4e-8 per base pair per generation (Zhu et al. 2018). For each species, we generated 1000 independent neutral genomic regions matching the length and SNP density of our empirical dataset. We applied SweeD v4.0 (Pavlidis et al. 2013) to each simulated dataset, using the same parameters as for the empirical data, to generate a distribution of maximum CLR values across 1,000 simulations, establishing an empirical null distribution.

Finally, we used SweeD to identify genomic regions under recent positive selection, using grid sizes of 10 kb and default parameters. To define coherent selective sweep regions from CLR peaks, we fitted cubic smoothing splines to the CLR scores along each chromosome using the *smooth.spline* function in R. Peaks exceeding 99.5% of the null distribution were considered significant. Outlier regions exceeding the significance threshold were merged when they were within 50 kb of each other (Tittes, Lorant, McGinty, et al. 2025).

### Comparison of selective sweeps among different species

After identifying sweep regions for each species, we performed pairwise comparisons to quantify shared sweeps among species. A sweep region was defined as shared if the overlapping genomic segment constituted >60% of the total region size in at least one of the species in the pair. To test whether the observed amount of sharing was statistically significant, we performed a permutation test. For each species, we generated a null distribution by creating 1,000 replicates of randomly sampled genomic windows that matched the number and size distribution of the observed sweeps. These random regions were sampled from non-overlapping portions of the genome to avoid bias from clustered sweep signals. For each permutation replicate, we calculated the total size of the overlap between these random windows from one species and the observed sweep regions from the other species. The statistical significance of the observed overlap was then assessed by calculating an empirical P-value, defined as the proportion of the 1,000 null replicates that yielded an overlap size greater than the observed overlap. Finally, to characterize the potential biological functions associated with signatures of selection, we performed Gene Ontology (GO) enrichment analysis on genes located within the identified selective sweep regions using the R package clusterProfiler v4.18 (Wu et al. 2021).

### Inferring modes of convergent adaptation

For sweep regions identified as shared between species pairs, we used rdmc (Lee and Coop 2017; Tittes 2020) to infer the most likely mode of convergent adaptation. This method compares the composite likelihoods of four distinct evolutionary scenarios: 1) neutral evolution, 2) independent selective sweeps acting on different mutations, 3) selection on shared standing genetic variation present in the ancestral population. 4) adaptive introgression (migration) of beneficial alleles between species. We followed the model configuration and parameterization settings adopted from previous study(Tittes, Lorant, McGinty, et al. 2025). Before model fitting, we expanded each identified sweep region by adding 10% of its total length to ensure sufficient flanking sequence for accurately estimating patterns of haplotype decay and recombination. Due to substantial differences in sweep region sizes across species pairs, we normalized SNP density by subsetting the total number of sites within each, maintaining a density between 1,000 and 100,000 SNPs per region. The recombination rate for each sweep region was approximated based on the genome-wide genetic map, assuming a recombination rate of 6.4e-8 (Zhu et al. 2018). Each shared sweep region was assigned to the evolutionary model with the highest log-composite likelihood.

To assess confidence in model assignments, we calculated the difference between the highest log-likelihood and the neutral model log-likelihood; regions with ΔCLE > 2.0 were considered to have support for a specific non-neutral evolutionary model. Comparisons between species divergence time and the number of shared sweeps were investigated with Pearson correlation coefficients. To account for the non-independence of species pairs due to shared evolutionary history, correlations were calculated using the Phylogenetic Generalized Linear Mixed Model (PGLMM) from MCMCglmm v2.36 package (Hadfield 2010).

## DATA AVAILABILITY

Newly generated Illumina whole-genome sequencing reads for 114 accessions are deposited in the NCBI SRA under the BioProject accession PRJNA1424665. Previously published whole-genome sequencing reads from 525 accessions can be accessed from the NCBI Sequence Read Archive (SRA) under the BioProject accessions PRJNA393611, PRJNA842753, PRJNA321480, PRJNA984685 and PRJCA009314. Geographic occurrence data were obtained from the Global Biodiversity Information Facility (https://www.gbif.org/). Environmental variables data are available from WorldClim (https://www.worldclim.org/).

## CODE AVAILABILITY

Analysis pipelines and customized scripts used in this study are available via GitHub at https://github.com/wang-tianpeng/Vitis_PopGen.

## ACKNOWLEDGMENTS

Rebecca Gaut generated the DNA and resequencing libraries, the UCI High Throughput Facility produced the resequencing data, and Edwin Solares provided technical assistance with high-performance computing. The work described herein was made possible by the Gaut lab’s GCluster high-performance computing resources as well as the resources of the Research Cyberinfrastructure Center at UC Irvine. This work was supported by the National Science Foundation USA, grants NSF DEB-2414478 and NSF DEB-2323123 to BSG and DC.

## Notes

### Competing Interest Statement

The authors have declared no competing interest.

