## Supplemental Figures for "The dynamics of introgression across an adaptive radiation: examining hybrid speciation and parallel adaptation in North American *Vitis*"

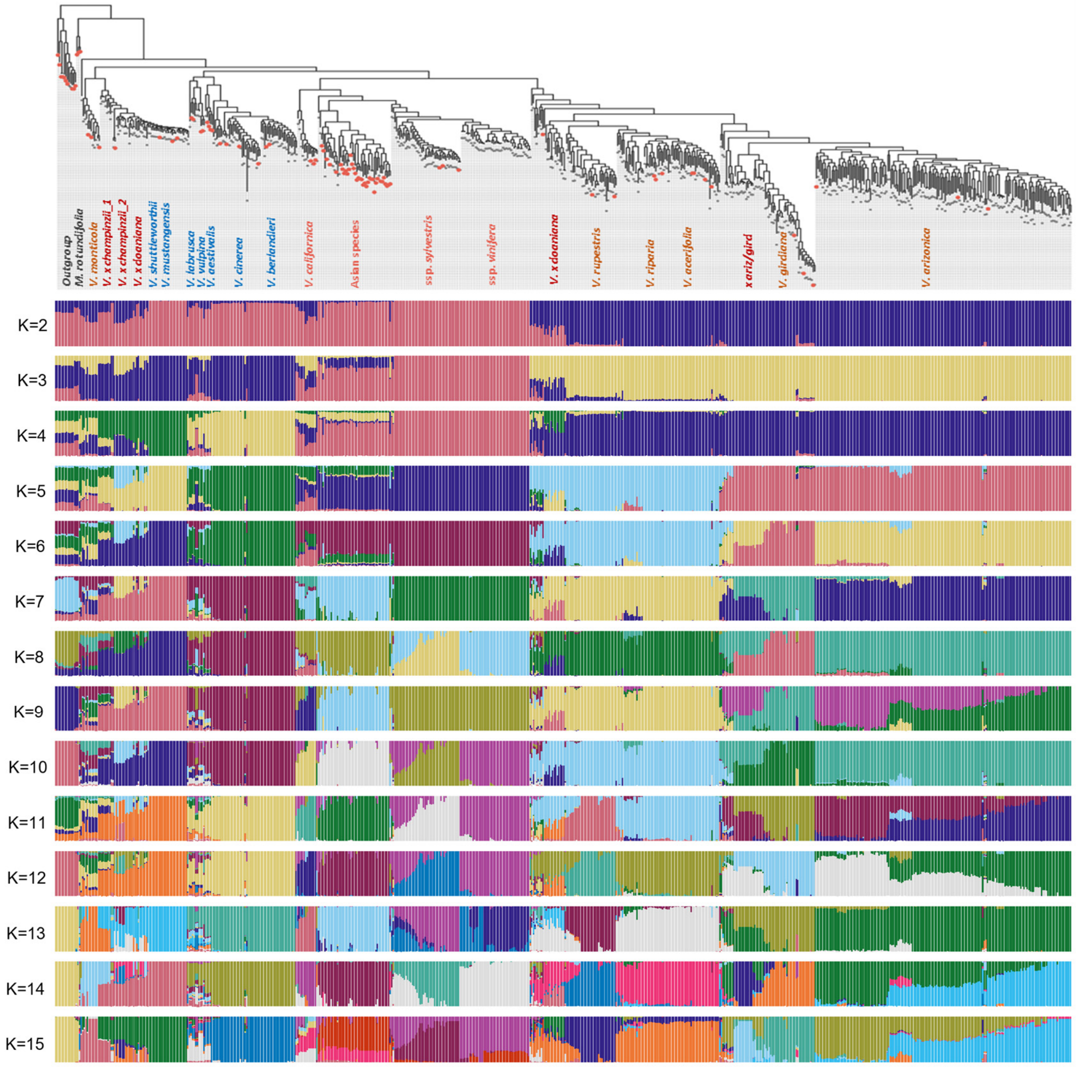


**Supplementary Fig. 1. Phylogeny and population structure of Vitis species. (A)** Maximum Likelihood tree of 639 Vitis accessions. Branch colors denote species. **(B)** Model-based Bayesian clustering performed with the number of ancestry kinship (K) from 2 to 15. The different colors of vertical bar represent contributions to the K-groups. Individuals are ordered based on their phylogenetic position.

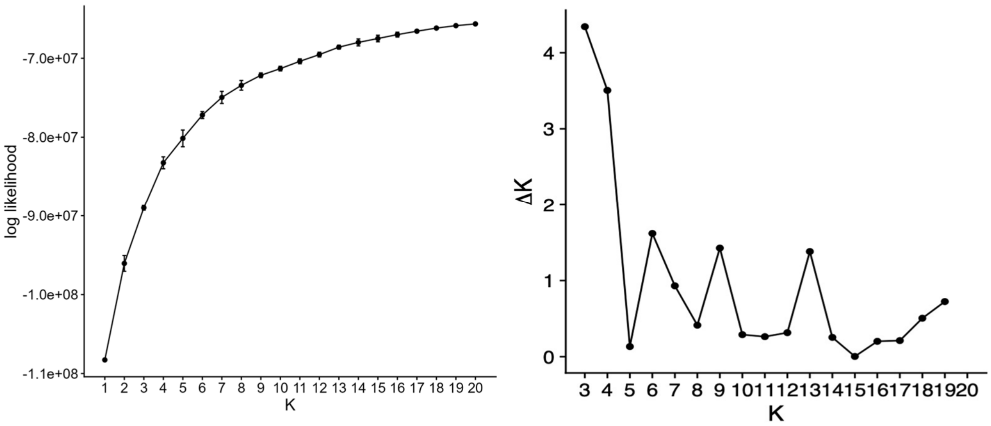


**Supplementary Fig. 2. Log-likelihood value for different population structure K value.**

Cross-validation errors for each deltaK from 3 to 19 are shown in the right panel.

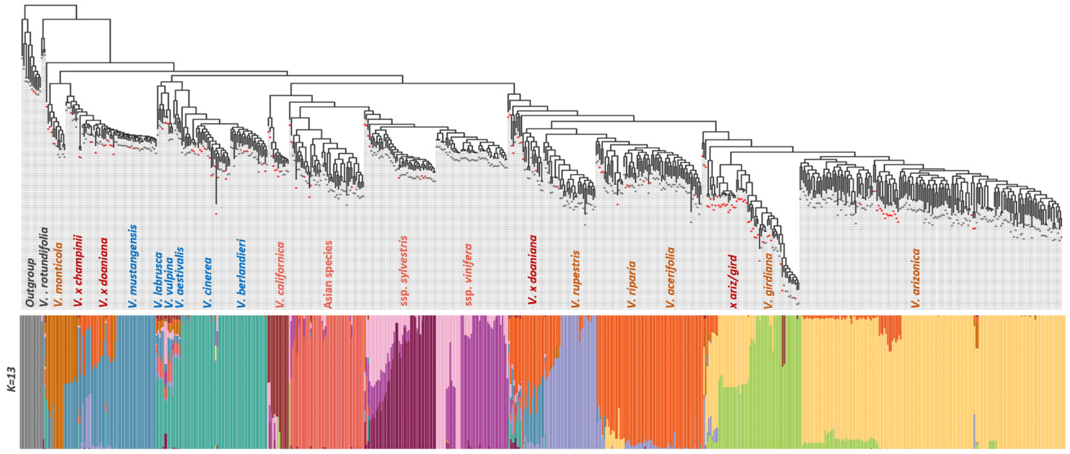


**Supplementary Fig. 3. Identification of putatively mislabeled or highly admixed individuals.** Individuals exhibiting substantial admixture (>20% ancestry from multiple clusters) or phylogenetic placement inconsistent with their assigned species were identified based on combined ancestry and phylogenetic analyses. Detailed information for all excluded individuals is provided in Table S2.

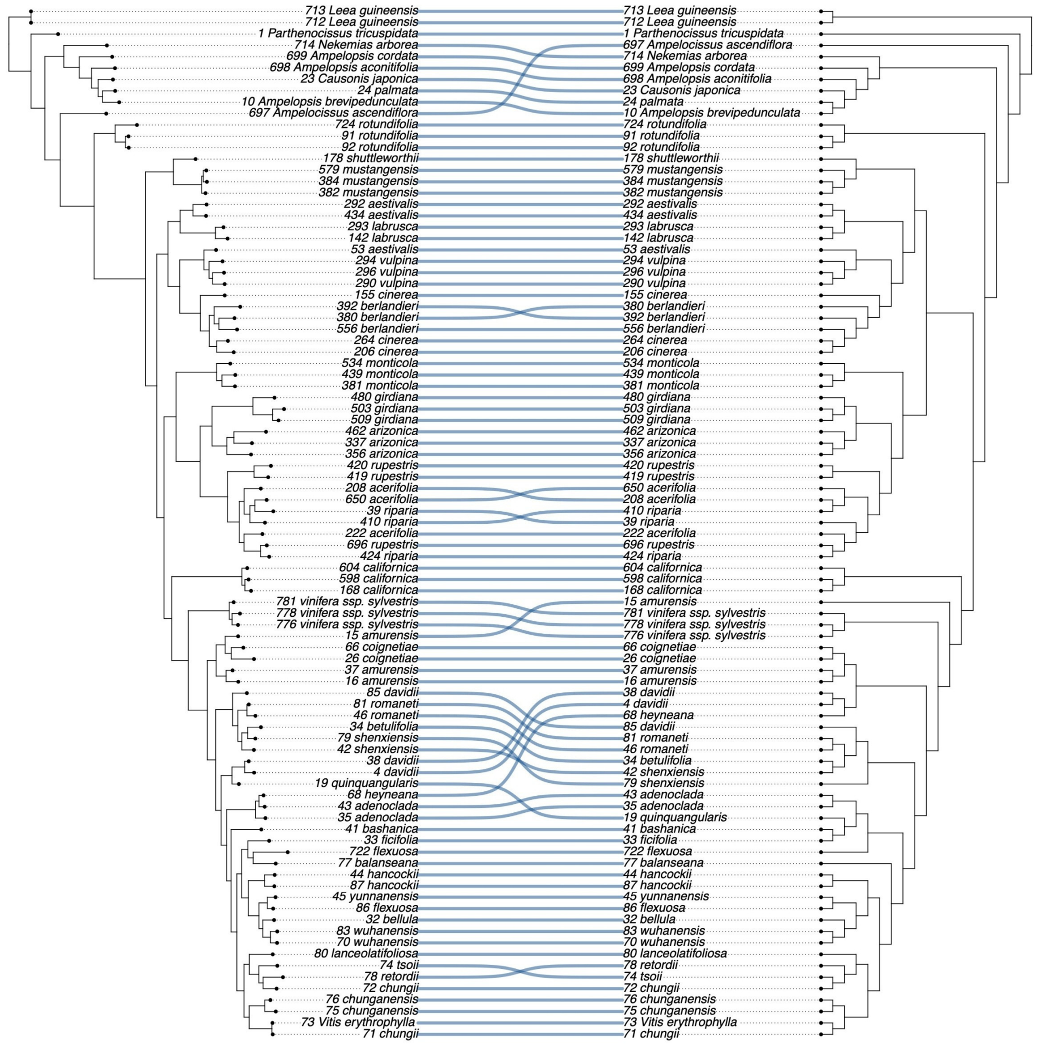


**Supplementary Fig. 4. Comparison of species phylogenetic topologies inferred by Raxml and SVD tree.** The tree on the left was inferred using RAxML (Maximum Likelihood), and the tree on the right was inferred using SVDquartets. Connecting lines link identical taxa between the two phylogenies to visualize topological concordance.

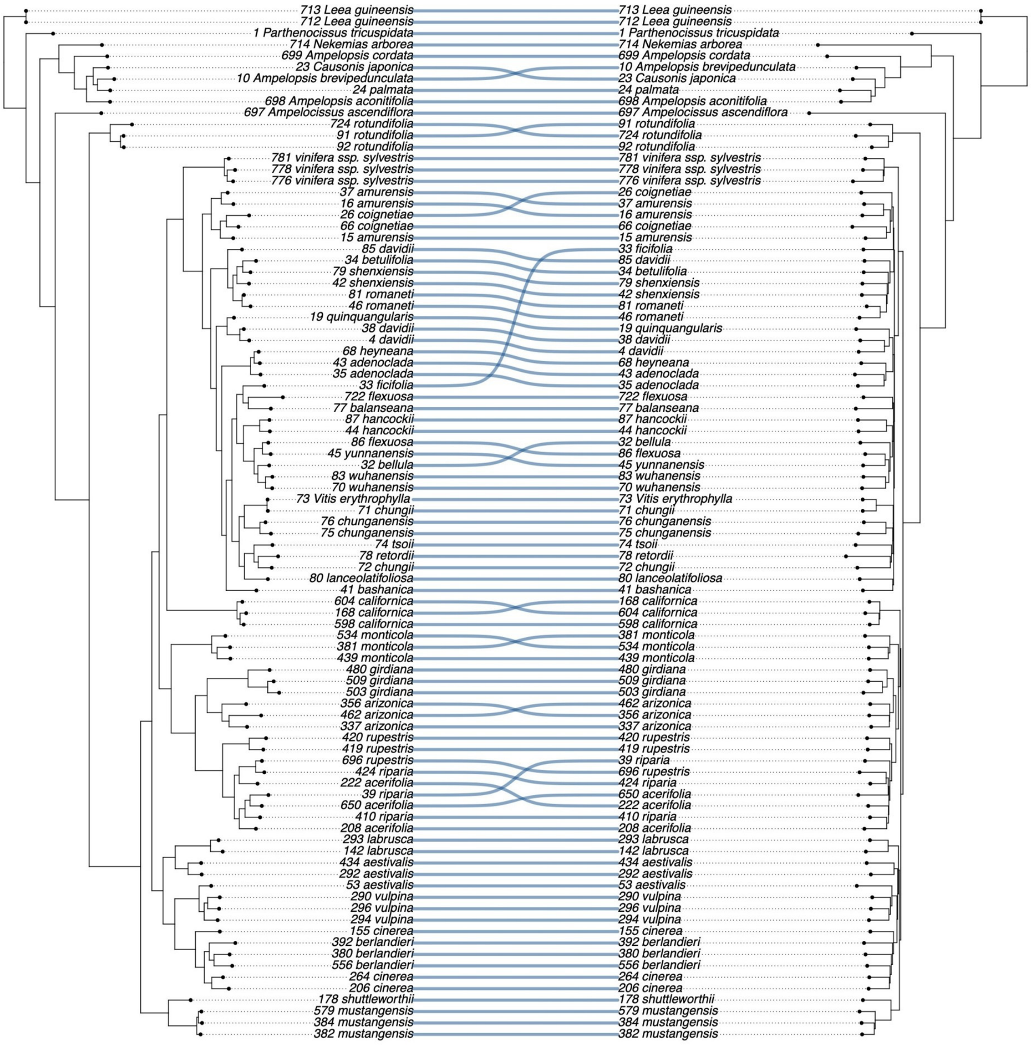


**Supplementary Fig. 5. Comparison of Raxml and kmer tree.** The figure displays the SNP-based RAxML tree on the left and the alignment-free K-mer tree on the right.

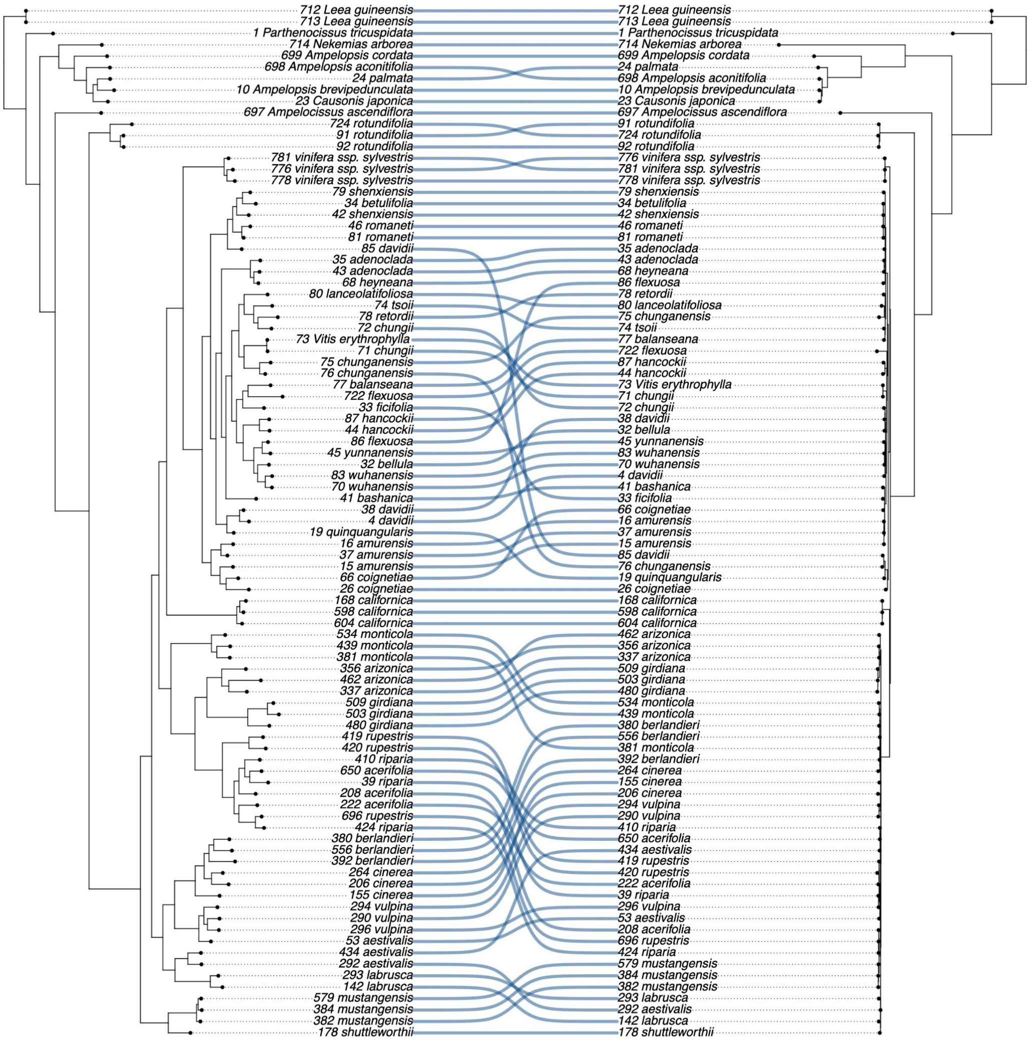


**Supplementary Fig. 6. Comparison of nuclear RAxML and plastid (Pt) trees.** The cophyloplot contrasts the nuclear SNP-based Maximum Likelihood tree (left) with the plastid haplotype-based tree (right). Connecting lines link corresponding taxa between the two phylogenies to illustrate differences in tree topology.

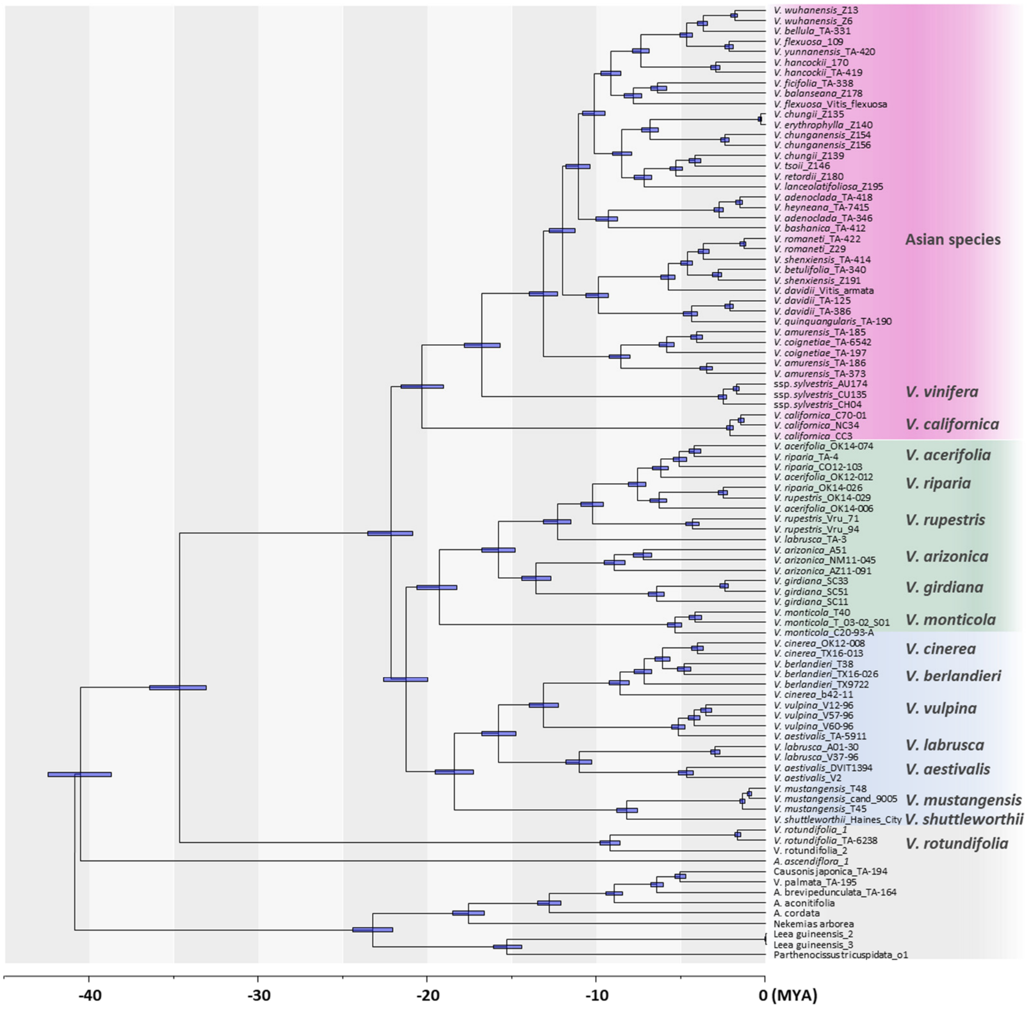


**Supplementary Fig. 7. Time-calibrated phylogeny of non-hybrid *Vitis* species estimated using BEAST2.** The chronogram was inferred using a relaxed molecular clock and a GTR nucleotide substitution model. Blue horizontal bars at the nodes represent the 95% Highest Posterior Density (HPD) intervals for the divergence time estimates. The scale axis at the bottom represents time in millions of years ago (Mya).

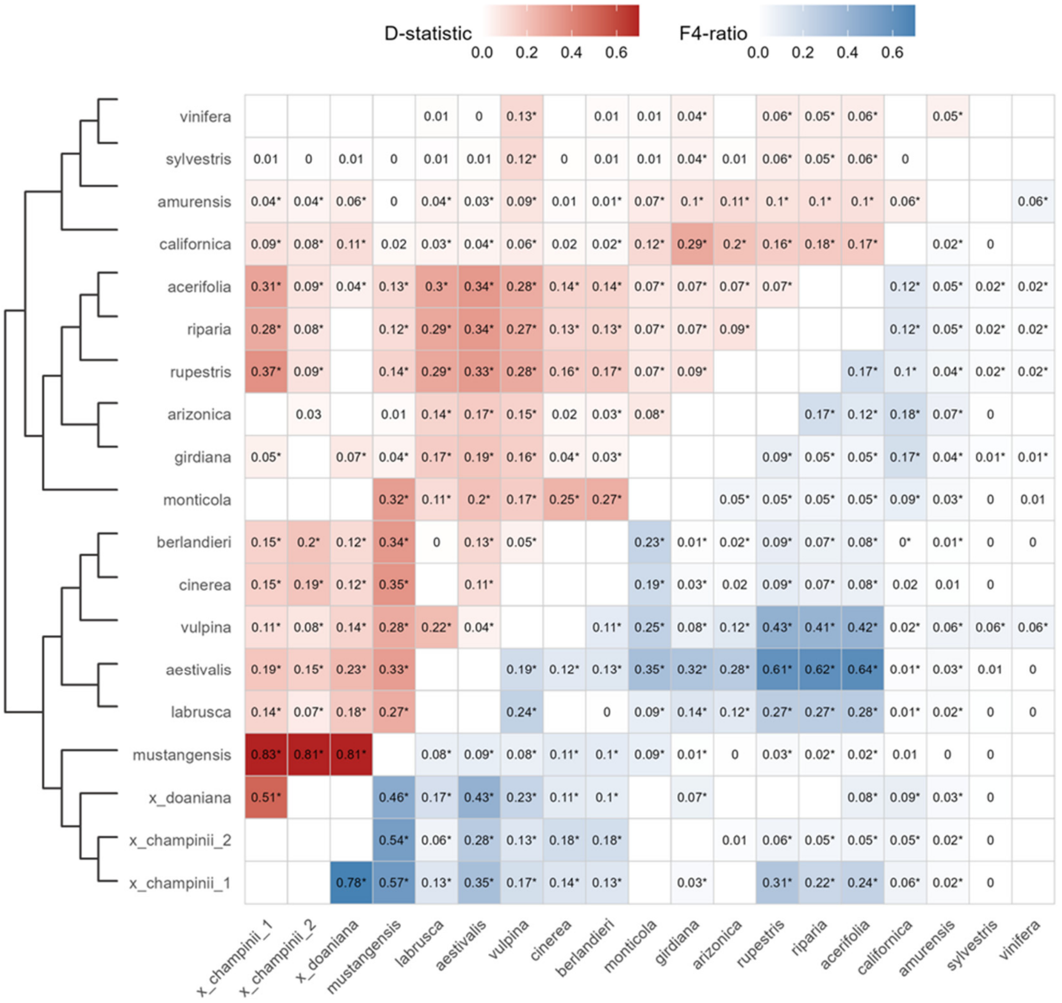


**Supplementary Fig. 8. Genome-wide introgression signals from pairwise D-statistics and F4-ratio**. Values marked with an asterisk denote significant introgression signals (Z-score > 4). Species are ordered according to the phylogeny shown on the left.

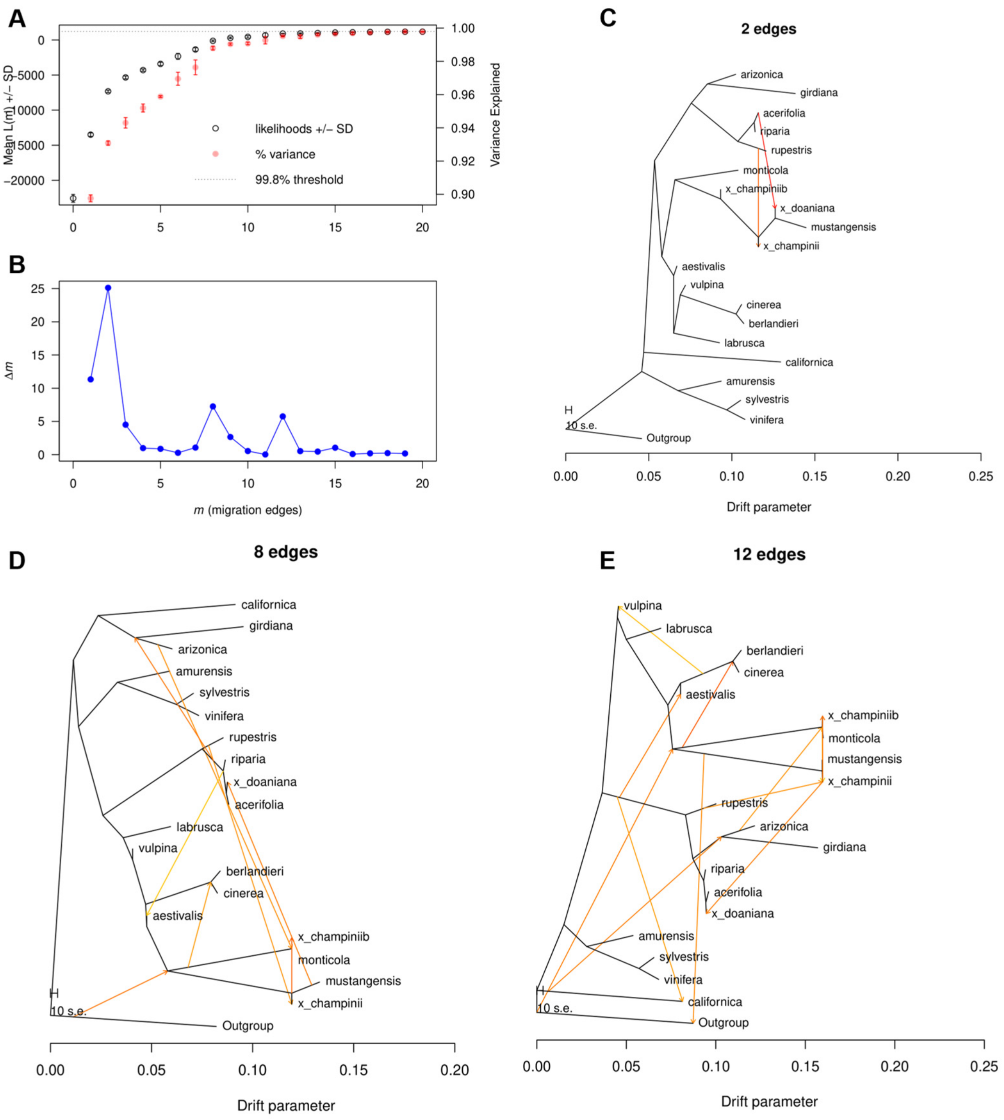


**Supplementary Fig. 9. Gene flow events among different Vitis species estimated by Treemix and OptM. (A)** The mean and standard deviation (SD) across 3 iterations for the composite likelihood (top axis) and proportion of variance (right axis) explained by models with 1-19 edges. **(B)** Distribution of deltaM statistic based on the second order rate of change in the likelihood with standard deviation considered. **(C-E)** Three models with different migration events that achieved >99.5% variance explained and the high deltaM values.

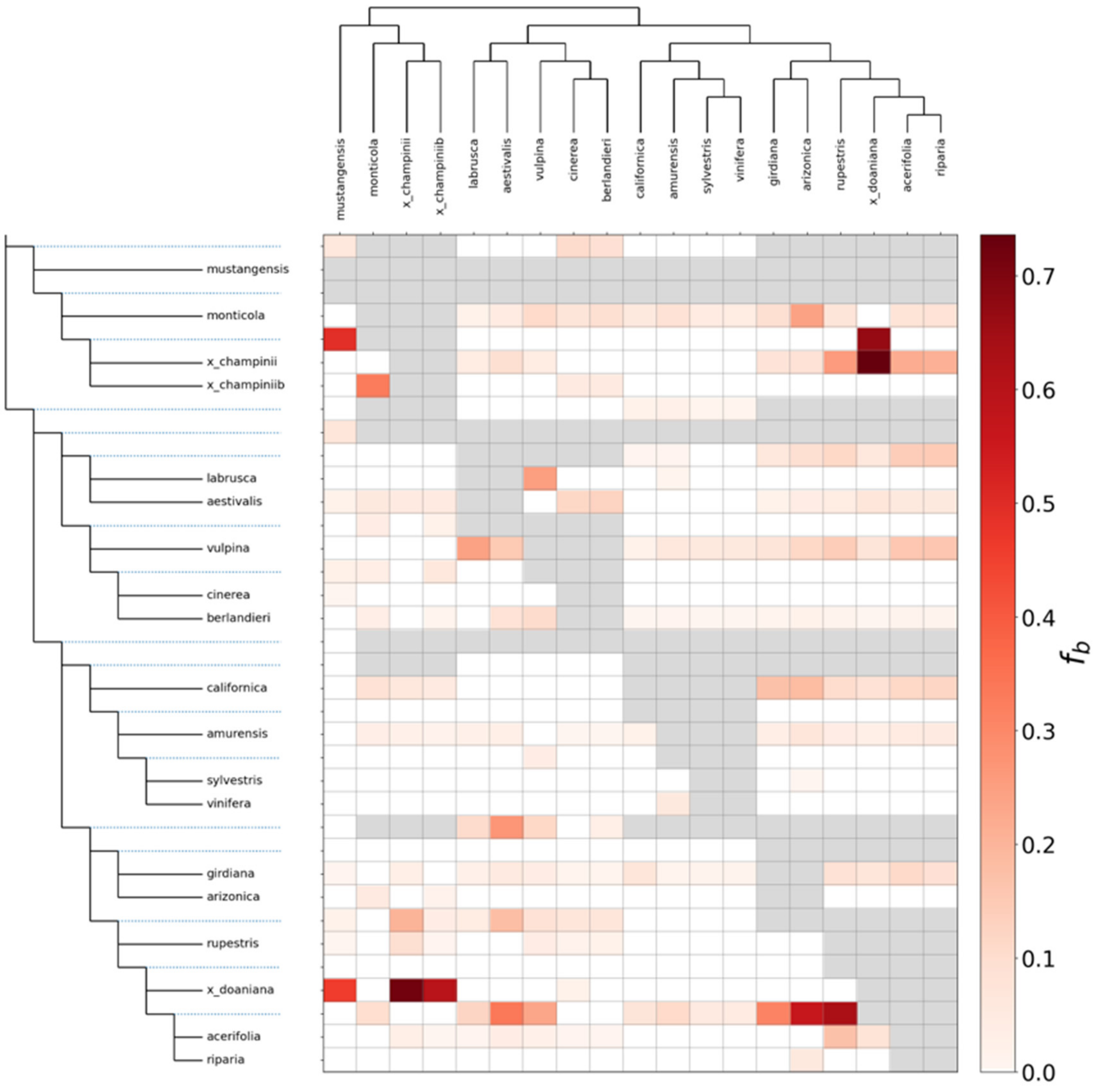


**Supplementary Fig. 10. Global signals of introgression inferred by the f-branch (fb) statistic.** The heatmap illustrates excess allele sharing between specific branches of the *Vitis* phylogeny (vertical axis) and potential donor species (horizontal axis). The phylogenetic tree on the left identifies the specific branch receiving the gene flow (indicated by dotted lines), while the tree on the top indicates the donor lineages.

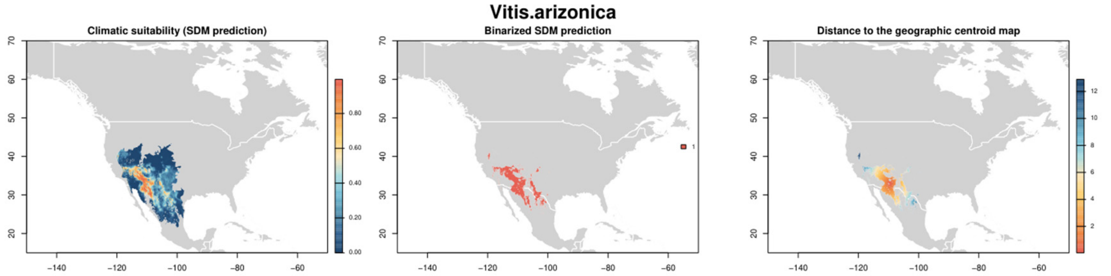


**Supplementary Fig. 11. (Additional file) Present Species Distribution Models (SDMs) for 14 North American Vitis species. (A)** Map displaying the probability of occurrence across the accessible area defined by biogeographic barriers (M) (Warmer colors indicate higher probability); **(B)** Binarized map derived from the SDM probability values using TSS and ROC thresholds; **(C)** Map showing for each location the distance to the geographic centroid of the binarized map (warmer colors show a lower distance to the geographic Centroid).

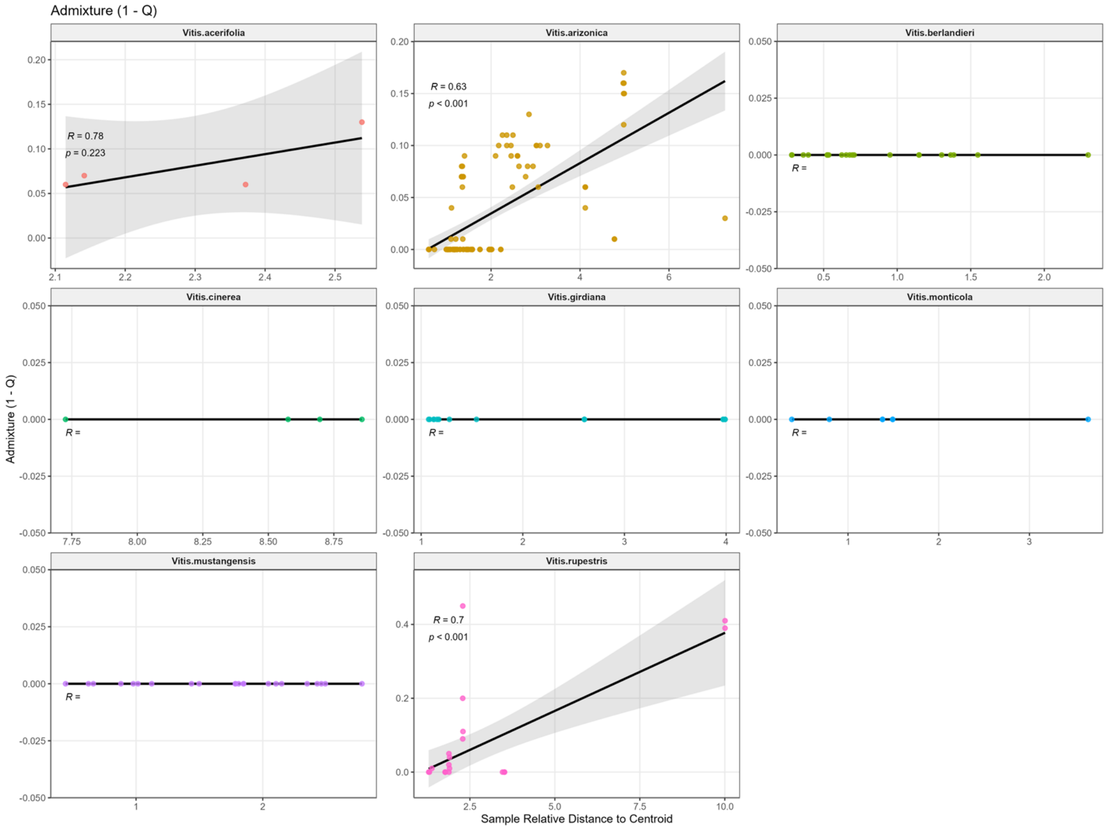


**Supplementary Fig. 12. Within-species Correlation between geographic distance to niche centroid and genomic admixture levels.** For each *Vitis* species, points represent sampled individuals plotted by their relative geographic distance to the species SDM niche centroid (x-axis) and their admixture level (y-axis). Admixture was quantified as 1−Q, where Q is the individual's maximum ancestry proportion inferred from the population structure analysis at K=13.

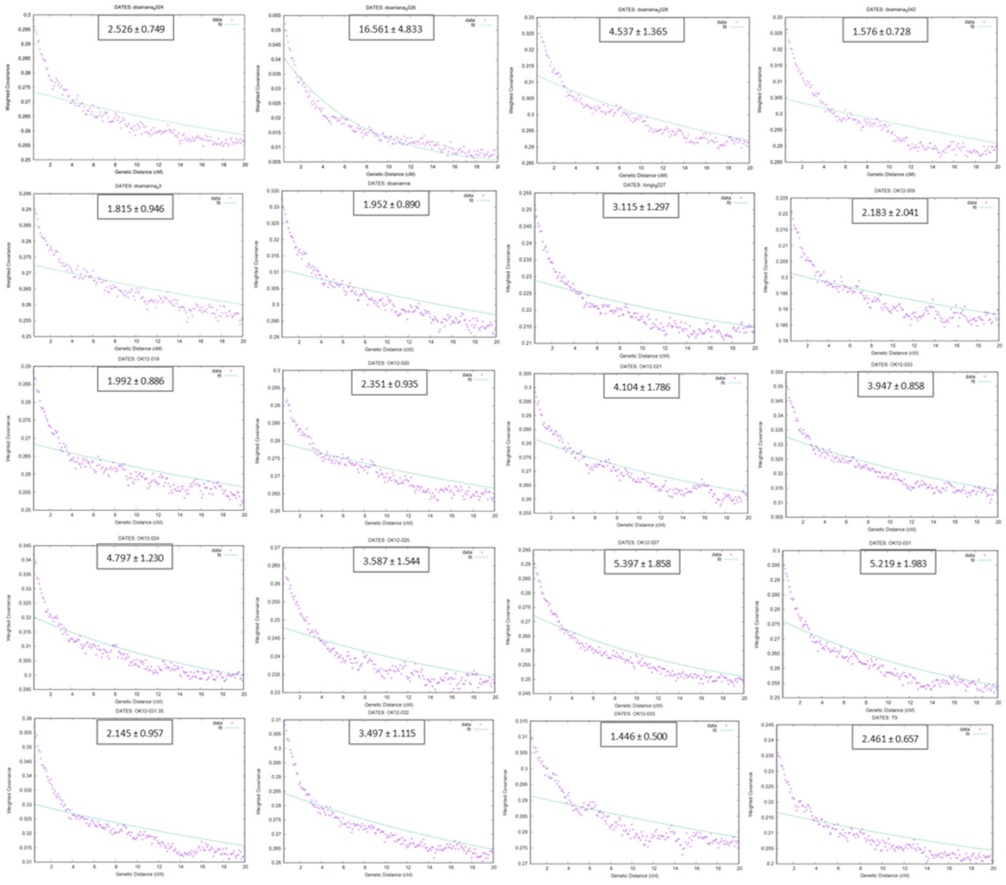


**Supplementary Fig. 13. Inference of hybridization timing for *V. doaniana* individuals using DATES.** The panels display the decay of weighted ancestry covariance (y-axis) as a function of genetic distance in centimorgans (cM, x-axis). The green line represents the fitted exponential decay curve used to estimate the time since admixture. The inferred number of generations since hybridization (mean +- standard error) is indicated below each plot.

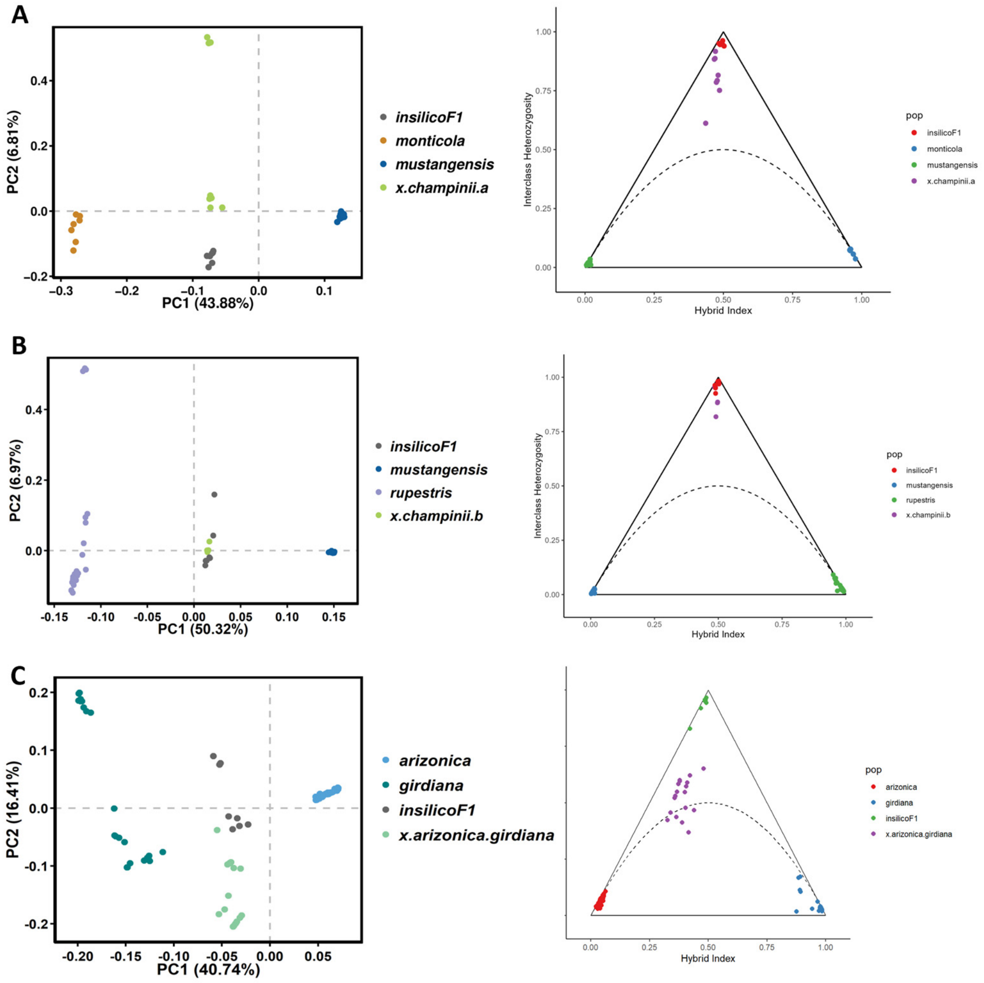


**Supplementary Fig. 14. Genomic evidence for recent hybridization in *V.* × *champinii* lineages and *V. arizonica/girdiana* hybrids.** For each putative hybrid group (rows), the left panel shows a principal component analysis (PCA) including the two inferred parental taxa and an in silico F1 control (constructed from parental genotypes). In all cases, putative hybrid individuals cluster near the in silico F1s and occupy intermediate positions between parental clusters along PC1. The right panel shows triangular (hybrid-index vs. interclass heterozygosity) plots. Putative hybrid individuals plot close to the apex and along the upper region of the triangle, consistent with very recent hybrid ancestry (F1/F2 and early backcross generations).

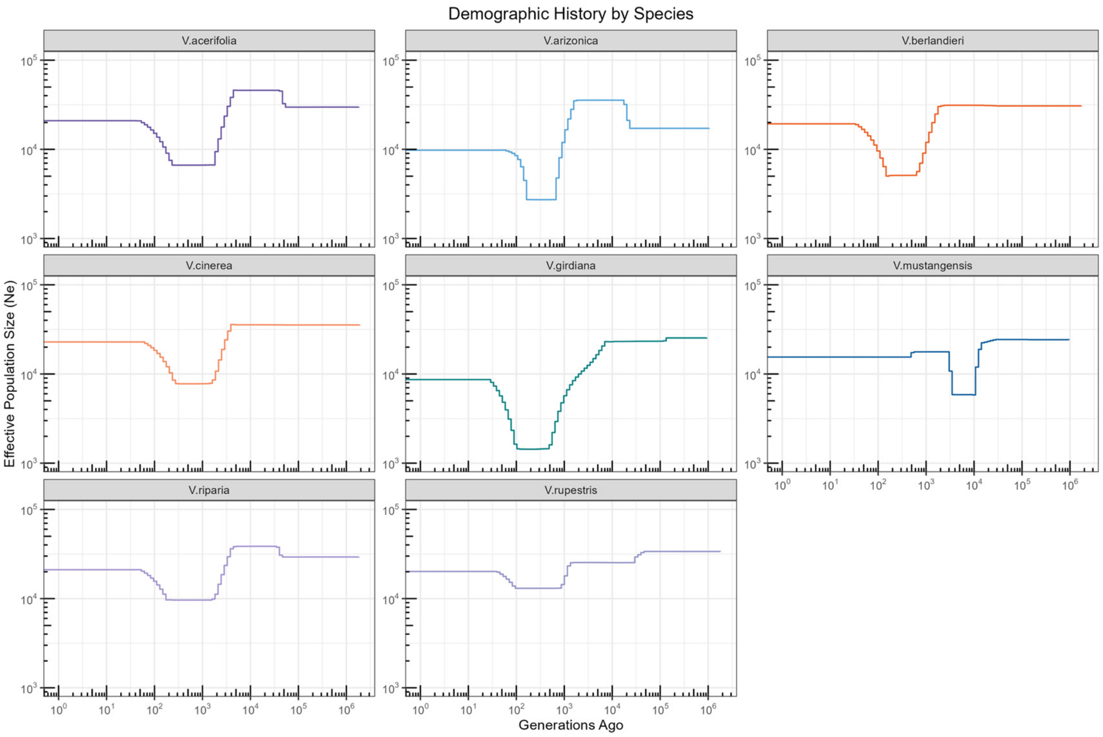


**Supplementary Fig. 15. Demographic history of each Vitis species inferred by mushi.** Generation estimates were inferred by assuming a generation time of 3 years and mutation rates of 5.4e-9 per site per generation, respectively. The panels display the changes in effective population size over time (generations ago) for selected Vitis species. The trajectories were reconstructed from the unfolded site frequency spectrum (SFS) using mushi v0.2. Both axes are plotted on a logarithmic scale.

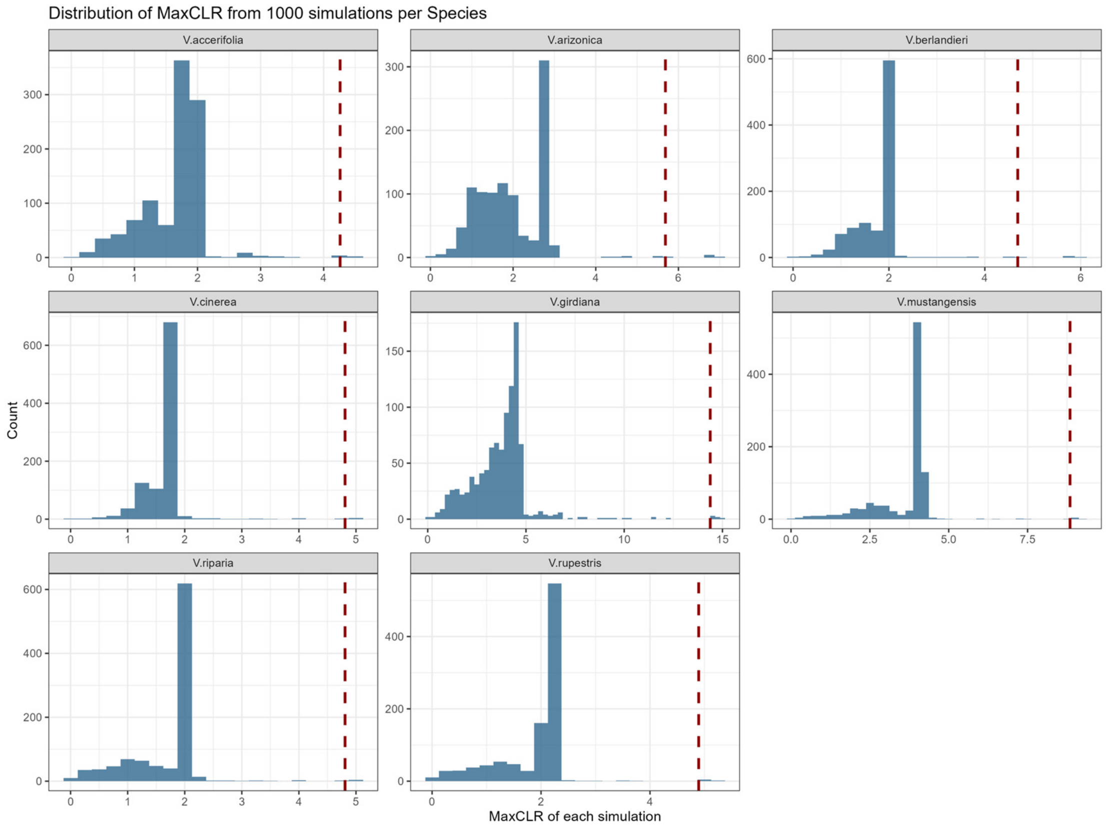


**Supplementary Fig. 16. Null distributions of the maximum Composite Likelihood Ratio (CLR) statistic derived from 1000 neutral coalescent simulations.** For each of the eight analyzed Vitis species, the histogram displays the distribution of the maximum CLR value obtained from 1,000 neutral simulations generated using msprime under estimated demographic histories. The vertical red dashed line marks the 99.5th percentile of each distribution, which served as the threshold for identifying selective sweeps in the empirical genomic data.

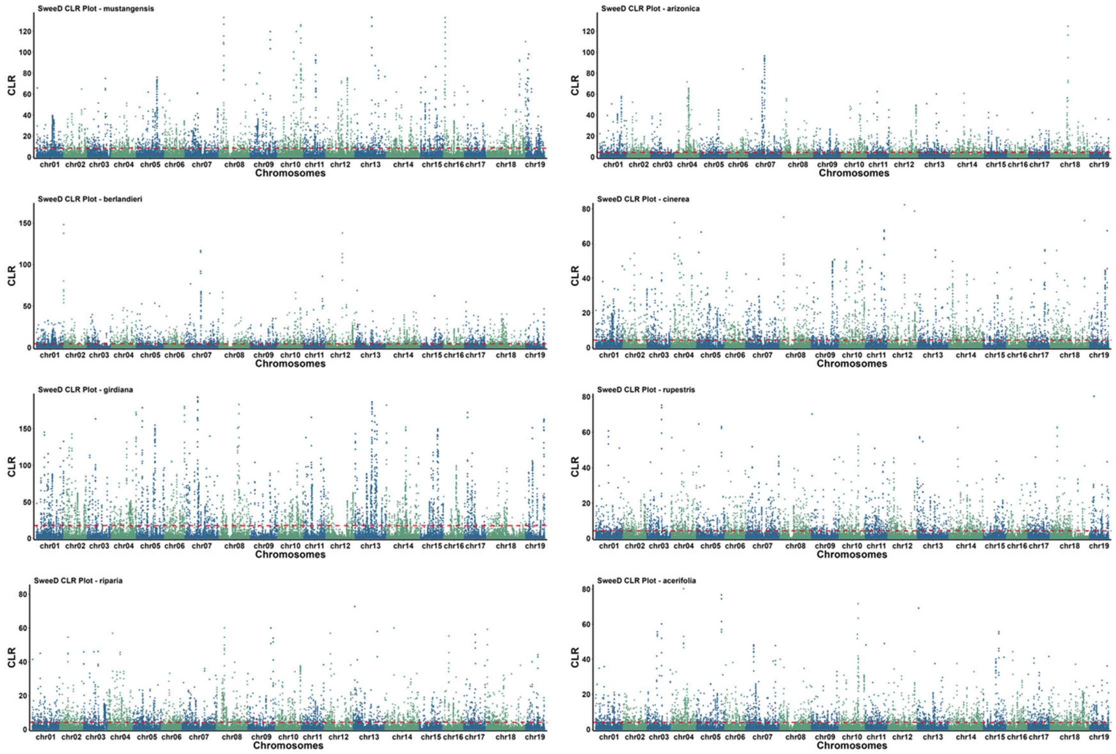


**Supplementary Fig. 17. Genome-wide scans for selective sweeps across eight North American *Vitis* species.** The Manhattan plots display the Composite Likelihood Ratio (CLR) statistic inferred by SweeD (y-axis) along the 19 chromosomes (x-axis).

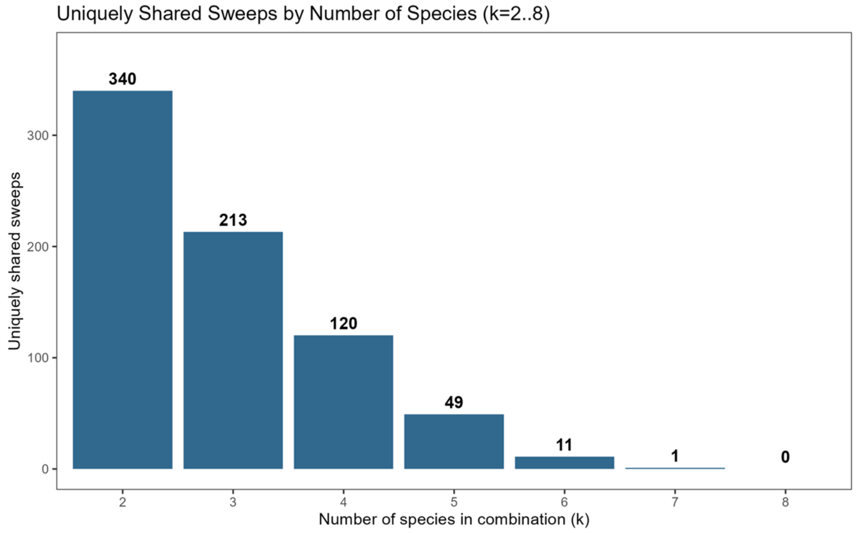


**Supplementary Fig. 18. Distribution of shared selective sweeps across multiple Vitis species.** The bar chart displays the number of selective sweeps shared by varying numbers of species, ranging from pairs (n=2) to sweeps shared across all eight analyzed species (n=8). A selective sweep was defined as shared if the identified genomic interval in one species overlapped by at least 60% of the sweep length in another species.

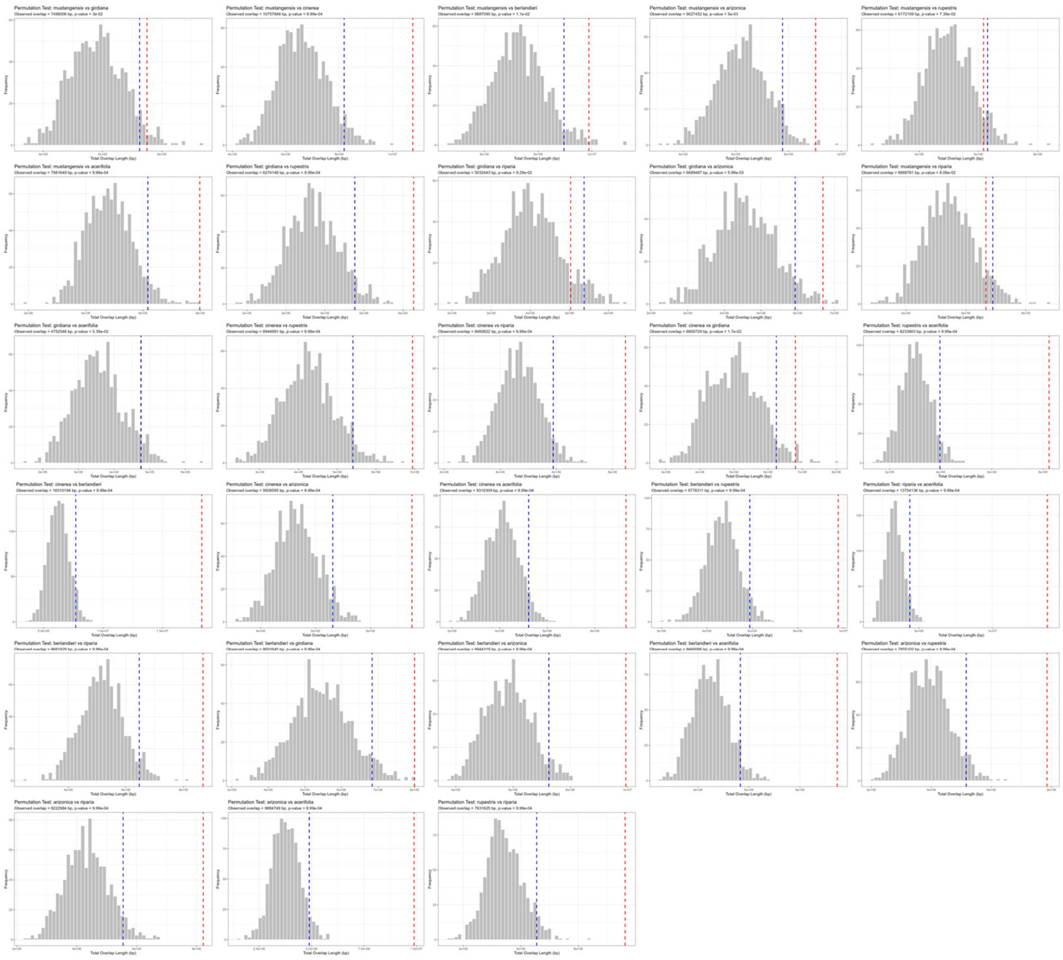

**Supplementary Fig. 19. Permutation tests for shared sweeps among Vitis species.** For each species pair, the histogram shows the null distribution of total sweep-overlap length (bp) expected under randomization. The red dashed line marks the observed total overlap length between the two species. The blue dashed line marks the 95th percentile of the null distribution. Observed values to the right of the blue line indicate significantly greater overlap than expected by chance.

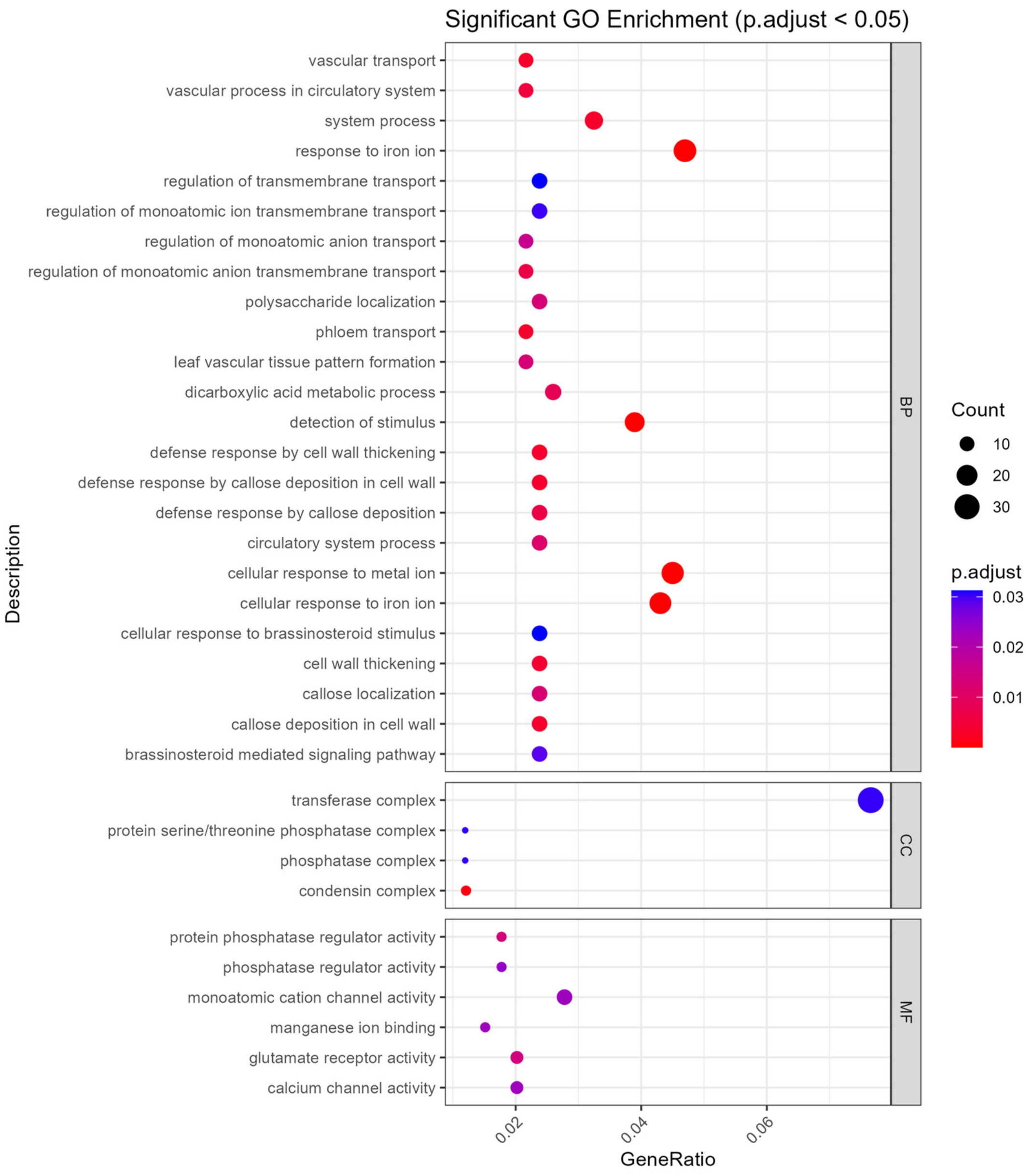


**Supplementary Fig. 20. Gene Ontology (GO) enrichment of selective sweeps among Vitis species.** The dot plot displays significantly enriched GO terms (P.adjust < 0.05) identified within selective sweep regions for eight Vitis species.
